# 48-spot single-molecule FRET setup with periodic acceptor excitation

**DOI:** 10.1101/156182

**Authors:** Antonino Ingargiola, Maya Segal, Angelo Gulinatti, Ivan Rech, Ivan Labanca, Piera Maccagnani, Massimo Ghioni, Shimon Weiss, Xavier Michalet

**Affiliations:** Department of Chemistry and Biochemistry, University of California Los Angeles, USA; Dipartimento di Elettronica, Informazione e Bioingegneria, Politecnico di Milano, Milan, Italy; Istituto per la Microelettronica e Microsistemi, IMM-CNR, Bologna, Italy.

**Author notes:** Electronic mail (Antonino Ingargiola). Electronic mail (Xavier Michalet).

**Keywords:** Single-molecule FRET, high-throughput, SPAD array

## Abstract

Single-molecule FRET (smFRET) allows measuring distances between donor and acceptor fluorophores on the 3-10 nm range. Solution-based smFRET allows measurement of binding-unbinding events or conformational changes of dye-labeled biomolecules without ensemble averaging and free from surface perturbations. When employing dual (or multi) laser excitation, smFRET allows resolving the number of fluorescent labels on each molecule, greatly enhancing the ability to study heterogeneous samples. A major drawback to solution-based smFRET is the low throughput, which renders repetitive measurements expensive and hinders the ability to study kinetic phenomena in real-time.

Here we demonstrate a high-throughput smFRET system which multiplexes acquisition by using 48 excitation spots and two 48-pixel SPAD array detectors. The system employs two excitation lasers allowing separation of species with one or two active fluorophores. The performance of the system is demonstrated on a set of doubly-labeled double-stranded DNA oligonucleotides with different distances between donor and acceptor dyes along the DNA duplex. We show that the acquisition time for accurate subpopulation identification is reduced from several minutes to seconds, opening the way to high-throughput screening applications and real-time kinetics studies of enzymatic reactions such as DNA transcription by bacterial RNA polymerase.

## I. INTRODUCTION

Detailed knowledge of the three-dimensional (3D) atomistic structure of macromolecular complexes is essential to understand their biological function. For decades, X-ray crystallography and nuclear magnetic resonance (NMR) spectroscopy have been the techniques of choice for obtaining atomically resolved macromolecular structures. More recently, single-particle cryo-electron microscopy (cryo-EM) has complemented these methods for determination of large macromolecular structures with the added ability to classify different conformations. However, macromolecules spontaneously and dynamically explore various conformations in equilibrium that are hard to capture by the above-mentioned methods. Understanding the functional roles of these structures requires a full dynamic picture. Single-molecule Förster resonance energy transfer (smFRET)^1^ has paved the way for studying such structural dynamics in biologically-relevant conditions. smFRET allows determination of each conformational state that may exist in an ensemble of macromolecular complexes as well as the distance between specific residues for each state^2–6^. Recently, several groups have implemented smFRET to measure distances between multiple different pairs of residues to construct 3D macromolecular structures of distinct conformations by triangulation and comparison with existing X-ray crystal structures^7–12^. Moreover, smFRET can measure the time-evolution of various distances between multiple FRET pairs, and hence report upon the dynamic 3D structure of a macromolecule undergoing conformational changes. Thus far, due to the requirement of low sample concentration imposed by the necessity to have no more than one molecule within the diffraction-limited confocal volume at a given time^1,13^, only very slow kinetics can be measured. Therefore, increased throughput is essential for both static and dynamic measurements of multiple distances.

To overcome this limitation, we recently introduced a multispot excitation scheme taking advantage of novel single-photon avalanche diode (SPAD) arrays^14–16^. We demonstrated that the resulting setup indeed allowed acquisition of single-molecule data comparable to that of standard single-spot setups, but with a throughput that scaled linearly with the number of excitation spots. We illustrated an application of this enhanced throughput by measuring the bubble closing kinetics during promoter escape in bacterial transcription^16^. While encouraging, these results were partially unsatisfactory because they were obtained with only 8 spots and also because the setup only incorporated a single laser, used for continuous excitation of the donor dye of the FRET pair^16^. Single-laser smFRET is unable to distinguish low FRET molecules (molecules with active donor (D) and acceptor (A) dyes in which the D-A distance is large compared to the Förster radius) from D-only molecules (*i.e.* molecules with a single donor, or dually-labeled molecules with an inactive or bleached acceptor). As the latter categories are present in most samples, it is important to identify and separate them from low FRET molecules of interest.

To address this problem, *Microsecond Alternated Laser EXcitation* smFRET (referred throughout as μsALEX for brevity), was introduced several years ago^17,18^, and later extended to pulsed laser excitation schemes (nsALEX^19^ or PIE^20^). Briefly, in μsALEX, two excitation lasers are alternated on and off every few tens of μs allowing separation of species with only a single active dye, *i.e.* D-only and A-only populations, from doubly-labeled species with both dyes active, the FRET populations. Indeed, only FRET populations emit a fluorescence signal during both D-excitation laser (due to excitation of the donor) and A-excitation laser (due to excitation of the acceptor). This, in turn, extends the number of FRET sub-populations that can be reliably identified within a sample in the regime of low mean FRET efficiencies, referred to throughout as “low FRET”. This scheme has since been extended to up to 4 laser excitations, allowing powerful molecular sorting applications^21,22^. A simplified version of this laser alternation principle was presented in ref. 23, where the D-excitation laser is left on at all times while the A-excitation laser is alternated. This Periodic Acceptor eXcitation single-molecule FRET technique^23^, referred to as PAX for brevity, simplifies the optical setup while maintaining the advantages of μsALEX, namely the ability to determine the number of D and A dyes in each detected molecule or to simplify the extraction of accurate FRET efficiency values^24^. Here we present a significant improvement to our original multispot setup by (i) introducing a 48-spot illumination and detection scheme, and (ii) implementing a 2-lasers, PAX illumination approach.

This paper is organized as follows. In Section II we briefly introduce the optical setup, the detectors (Section II A) and the modulation scheme (Section II B). In Section III we report single-molecule measurements, starting with a brief description of samples (Section III A) and data analysis (Section III B). To demonstrate the uniformity across spots, we study the burst peak photon rate (Section III C) and E-S histograms (Section III D). Finally, we compare single-spot μsALEX and 48-spot PAX measurements (Section III E). We conclude with a brief summary and perspective in Section IV.

### A. Software and data availability

Software used to operate the multispot setup (including LCOS-SLM pattern generation, LabVIEW-FPGA time-stamping code, piezo-motor control, etc.) is provided in various repositories available on GitHub^25^. Software used for data analysis can be found in the 48-spot-smFRET-PAX-analysis repository^26^. Links to specific analysis notebooks are added in the caption of each figure. Data files are publicly available on Figshare^27^.

## II. SETUP DESCRIPTION

In this Section, we provide a brief description of the setup. A more detailed description can be found in Appendix A, while details of laser and SPAD array alignment can be found in Appendix B and C.

A schematic of the setup is shown in Fig. 1. The setup includes two 1 W CW excitation lasers (green: 532 nm, red: 628 nm) where only the red laser is modulated via an acousto-optic modulator (AOM). After polarization adjustment and beam expansion, the two lasers are phase modulated by their respective LCOS-SLM, generating two 48-spot patterns on an image plane in front of each LCOS-SLM (*LCOS image plane*). The two modulated laser beams are then combined by a dichroic mirror (*DM*_*MIX*_) and recollimated (*L*_3_) before being focused into the sample by a high numerical aperture (NA) water immersion objective lens (60X, NA = 1.2, Olympus, Waltham, MA). Emitted fluorescence is collected by the objective lens, separated from the excitation wavelengths by a dual-band polychroic mirror (*DM*_*EX*_), and focused by a tube lens (*L*_2_) into the microscope’s bottom image plane. Next, emitted fluorescence light is recollimated (*L*_4_), separated into donor and acceptor spectral bands by a dichroic mirror (*DM*_*EM*_), and focused into two different 48-pixel SPAD arrays mounted on motorized micro-positioning stages (xyz vectors). The system is aligned such that each SPAD is optically conjugated to one excitation spot in the sample.

Output from the detectors (one TTL pulse train per SPAD) is processed by a field-programmable gate array (FPGA) equipped board (PXI-7813R, National Instruments, Austin, TX) that performs photon time-stamping with 12.5 ns resolution and transfers data asynchronously to the host PC. The host PC runs a LabVIEW acquisition software, which displays the binned signal recorded from all 96-channels in real-time as 96 color-coded time traces, implements alignment routines, and saves the data to disk. After data acquisition, file conversion to the Photon-HDF5 format^28^ and analysis is performed on a second PC, therefore allowing non-stop acquisition of sequential files.

### A. Detectors

The current 48-spot setup employs two identical 12x4-pixel SPAD arrays whose architecture and performance has been previously presented^29^. Here we describe only their most relevant features. Each SPAD has a 50 μm diameter active area, the array being comprised of 4 rows of 12 pixels (4x12) separated by 500 μm in both directions.

To easily integrate the detectors into the setup, we developed a photon-counting module that integrates a 48-pixel SPAD array and the electronics required for device operation, data acquisition, and transfer. The SPAD array is housed into a hermetically sealed chamber separated from the rest of the module by O-rings and uses a thin glass plate as an entrance window. The chamber is regularly flushed and filled with dry nitrogen gas to prevent condensation, making it possible to mount the array on a double-stage Peltier element to cool the detector down to temperatures of approximately −15°C. At this temperature, the dark count rate (DCR) is significantly reduced, thus increasing the signal-to-background ratio of the instrument. Photon-counting pulses are transferred through a standard SCSI connector. This allows for easy connection of the module to general-purpose data acquisition, or breakout adapter boards when different connectors or pulse shapes are required. Alternatively, an on-board FPGA (Spartan 6 SLX150, Xilinx, San Jose, CA) can be used to time-stamp counts detected in each of the 48-channels with a time resolution of 10 ns. This information is then sent to the host PC via a high-speed USB link. A C-mount thread around the entrance window of the photon-counting module allows for easy and reliable connections to the optical setup.

**Figure 1.**
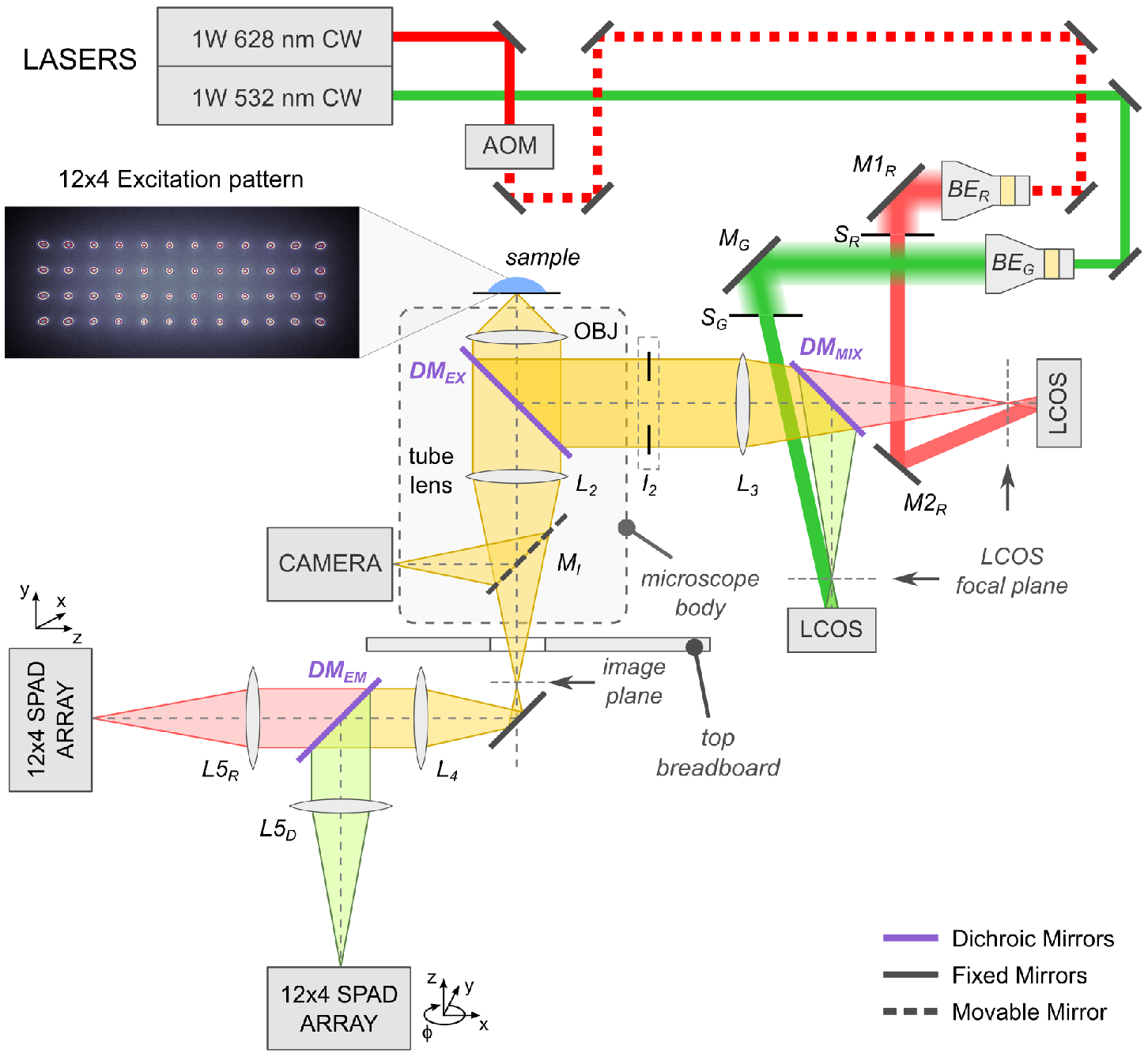
Schematic of the 48-spot PAX setup. The lasers and detectors are on the main optical table, while the microscope (enclosed in the dashed box), beam expanders, and LCOS-SLMs are located on a raised breadboard. See main text and Appendix A for a detailed description.

The two SPAD arrays used in the current 48-spot PAX setup are operated at a temperature of −10°C. The photon detection efficiency (PDE) reaches a maximum of ~45% at 550-580 nm (donor dye, ATTO550 emission peak) and drops to ~30% at 670 nm (acceptor dye, ATTO647N emission peak)^29,30^. The PDE is highly uniform over the array, with a peak-to-peak spread of only a few percent^29^. Fig. 2 shows DCRs for the two 12×4 SPAD arrays. Approximately 80% of the pixels have a DCR lower than 1,000 counts per second (cps), and the worst performing pixel has a fairly high DCR of nearly 6 kcps.

The 48-spot PAX results were compared to those of a state-of-the-art single-spot μsALEX setup previously described in 16. The single-pixel SPADs (SPCM-AQRH, Excelitas Technology Corp., Waltham, MA) used in the μsALEX setup are characterized by a PDE of ~60% at 550 nm and ~70% at 670 nm, with notably better sensitivity in the donor emission band and a PDE that is more than twice a high for the acceptor emission band. For this reason, the μsALEX setup is expected to be at least twice as sensitive in the A-channel than the 48-spot setup. A detailed comparison of the different SPAD technologies for single-molecule measurements is reported in 30.

### B. 48-spot pattern

The 48 excitation spots are generated independently for each wavelength by phase modulation of the incoming laser wavefront, as previously described in 16 and 32. The phase modulation operates in direct space rather than Fourier space and implements the phase profile of a Fresnel lenslet array. Similar direct-space modulation using a different spatial arrangement of the phase pattern on the LCOS-SLM have also been demonstrated for multi-confocal fluorescence correlation spectroscopy (FCS)^33^.

Fig. 3 shows the emission pattern from a high-concentration dye sample upon green (Panel A) and red (Panel B) laser excitation, as seen by a camera mounted on the microscope side-port. The two patterns are aligned to maximize overlap of each of the 48 spots.

**Figure 2.**
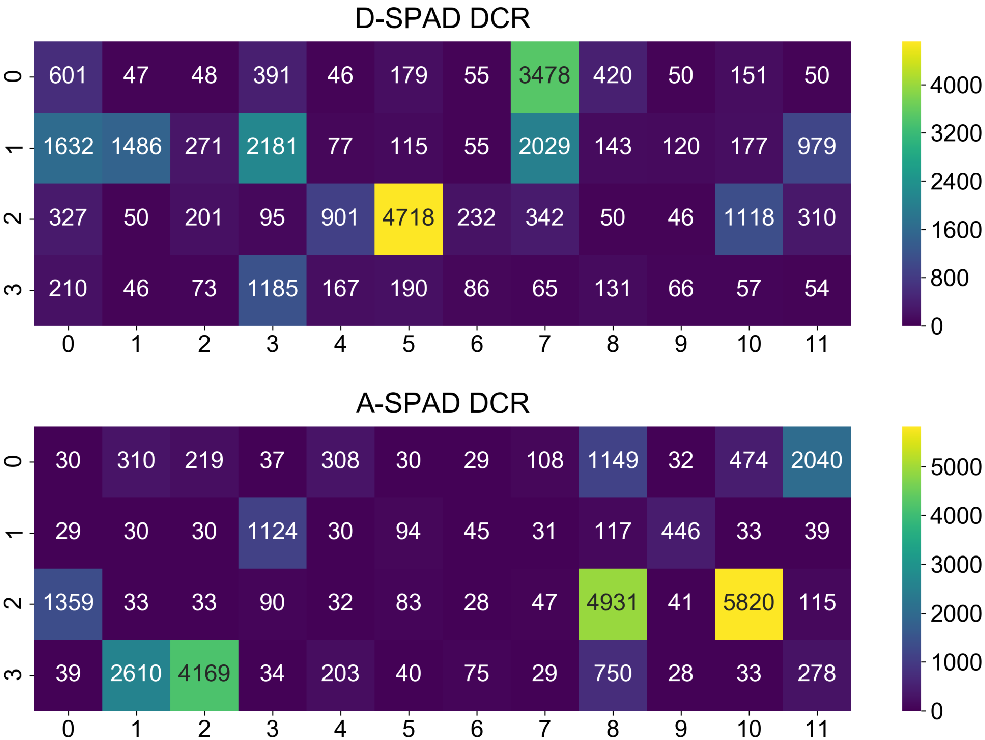
Heatmaps of DCRs for the 12x4 D- and A-SPAD arrays used in the 48-spot PAX setup. DCR values in counts per second (cps) are indicated in each pixel. More details and data can be found in the accompanying DCR analysis notebook^31^.

**Figure 3.**
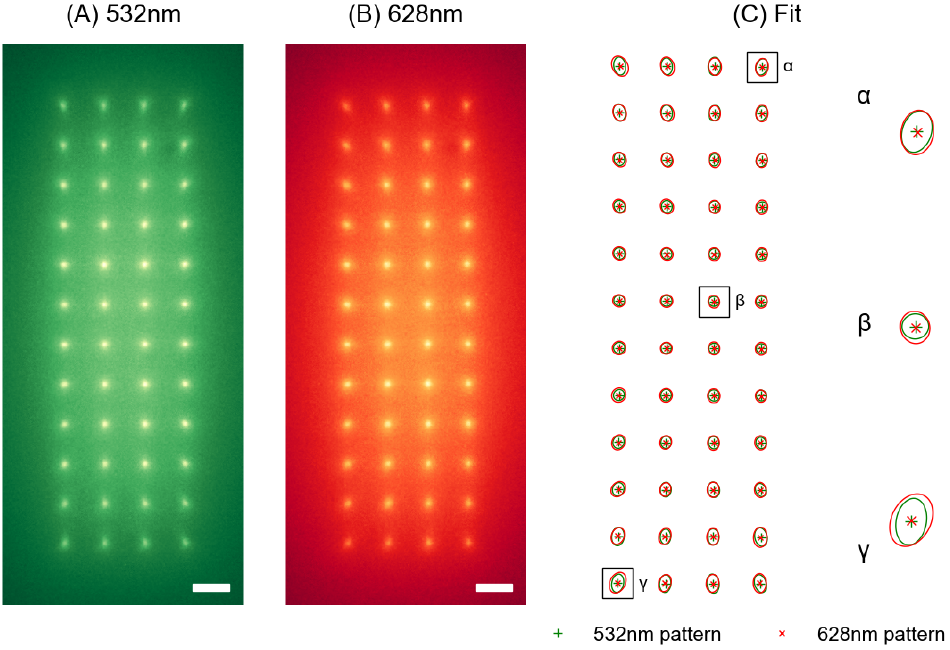
The 12x4 multispot pattern for green (A) and red
(B) excitation and Gaussian fit of the spots (C). The pattern is acquired by a camera mounted on the microscope side port (see Fig. 1) using a solution of ATTO550 and ATTO647N dyes at high concentration (~100 nM). Fluorescent images obtained upon 532 nm or 628 nm laser excitation were acquired separately and are reported in green and red intensity levels in panels (A) and (B), respectively. Scale bars are 5 μm. To assess the alignment, each spot in the two images is fitted with a 2D Gaussian function. Panel (C) reports an overlay of the fitted peak positions and a contour of the Gaussian waist for 532 nm (*green*) and 628 nm (*red*) images. A closer look of 3 representative spots is reported on the right. The elliptical shape and tilt of the Gaussian is due to geometrical aberrations. More details can be found in the accompanying alignment notebook^34^.

Overlap of the two wavelengths and centering with respect to the optical axis is assessed by 2D Gaussian fitting of each individual spot as reported in Panel C. Full details on the alignment procedure and pattern assessment can be found in Appendix A 1.

## III. SMFRET MEASUREMENTS

### A. Samples

Single-molecule measurements were performed with 40 base-pair (bp) double-stranded DNA (dsDNA) molecules labeled with ATTO550 (D) and ATTO647N (A) dyes (ATTO-TEC GmbH, Siegen, Germany) attached to different DNA bases, yielding different inter-dye distances.

D-A separation of 12 bp and 22 bp were used in these experiments, as they cover the typical range of distances that can be accurately measured with smFRET using this dye pair. Samples were diluted to single-molecule concentration (~50 pM) in TE50 (10 mM Tris pH 8.0, 1 mM EDTA, and 50 mM NaCl) or in “transcription buffer” (40 mM HEPES-NaOH pH 7, 50 mM KCl, 10 mM MgCl_2_, 1 mM DTT, 1 mM MEA, BSA: 100 μg/ml)^35^. TE50 buffer was used for all measurements excluding data collected for Fig. 12 and 13. The transcription buffer reduces photo-bleaching, and it was necessary so that a single sample could be measured consecutively on both setups without significant loss of fluorescence. Full details regarding the DNA samples are provided in ref. 16.

### B. Analysis

We analyzed data using standard μsALEX methods^24^ with modifications required for PAX^23^. The three analysis steps include: (a) background estimation, (b) burst search and (c) burst selection. Background estimation, which is needed to correct the burst counts in the different photon streams, was performed over 10 s time windows in order to account for possible background variations during the measurement. Burst searches were performed independently for each spot using the sliding-window algorithm^13^ and a constant-rate threshold for all spots^36^. Burst selection is performed taking bursts with size larger than a specified threshold, where burst size is either defined according to eq. D5 or D14. To isolate the FRET populations, we additionally filter bursts with *DA*_*ex*_*A*_*em*_ counts larger than a second specified threshold. Full numerical details can be found in the relevant notebook^26^.

The main result of the μsALEX and PAX analysis methods is a so-called E-S two-dimensional histogram, where each burst is represented by a pair of values (*E*, *S*) computed from the distinct photon stream intensities (see Section III D). The E-axis in that histogram can repre-sent either the FRET efficiency or, more commonly, the uncorrected FRET efficiency *E*_*PR*_, known as proximity ratio. *E*_*PR*_ is easier to compute than *E* and provides a suitable approximation for identifying sub-populations. However, while it is not the objective of this study, when the purpose is extracting D-A distances, all the correction coefficients need to be accurately estimated in order to compute *E. S*, or “stoichiometry ratio”, is a quantity which typically has a value ~0:5 for doubly-labeled, ~0 for A-only, and ~1 for D-only species. D and A-only species regions in the E-S histogram also include doubly-labeled molecules with one inactive dye due to photo-blinking or bleaching. Unlike the corrected stoichiometry ratio *S*_*γβ*_ (eq. D9), the uncorrected ratio *S* (eq. D8) can exhibit a dependence on *E* and for doubly-labeled molecules is not necessarily centered about 0.5. The use of the (*E*, *S*) pair (corrected or uncorrected) allows separation of singly and doubly-labeled species and distinguishing FRET sub-populations within the doubly-labeled population. Full definitions of *E* and *S* as well as comparisons between ALEX and PAX variants are reported in Appendix D.

In this paper, we report proximity ratios *E*_*PR*_ com-puted according to eq. D6, and a “modified stoichiometry ratio” *S*_*u*_ defined in eq. D19. *S*_*u*_ is a variant of the classical PAX stoichiometry ratio^23^, which reduces the effect of shot noise and improves the separability of D-only and FRET populations. More details on *S*_*u*_ can be found in Appendix D 1. Note that throughout this work, the results of the two leftmost spots in the second row are missing because of an active quenching circuit (AQC) failure in the D-SPAD array.

### C. Peak photon rate

The peak photon rate reached in each burst reports on the peak PSF intensity^16^. Fig. 4A shows the background-corrected peak photon rate distributions with their characteristic exponential tails. Fig. 4B-E show, for different photon streams, heatmaps of the peak photon rate mean values, *i.e.* the decay constant of the exponential tail. Due to the Gaussian profile of the excitation beam and to geometric aberrations, the lateral pixels receive a lower signal intensity than the central pixels. As a result, the peak photon rate decreases and fewer single-molecule bursts are detected in the lateral spots. Despite this decrease in excitation intensity, the positions of the *E*_*PR*_ and *S* peaks remains quite uniform across the spots (see Fig. 7 and 9). An exception can be seen in Fig. 4C,E, where the pixel at position (1, 7) in the A-SPAD array detects fewer photons than its neighbors, an effect we ascribe to a lower PDE of that pixel, possibly due to a lower applied overvoltage. For this spot (19), we observe a noticeable bias in *E*_*PR*_ and S quantities (see Fig. 7 and 9).

By comparison, Fig. 5 shows the distribution of peak photon rates obtained with the μsALEX setup. The absolute power delivered in each spot in the multi-spot setup is difficult to measure. Therefore, we determined an optimal D-laser power (200 mW at laser output) so that the peak photon rate distribution in the central spots was comparable to the single-spot peak photon rate (190 μW measured before the objective). The A-laser power was set to 400 mW (laser output before the AOM), as a trade-off between the need to compensate for the lower PDE in the A-channel and limiting sample photo-bleaching and thermal instabilities.

Comparing Figs. 4A and 5 it is clear that reduced sensitivity in the A-SPAD array results in lower peak photon rates in the *D*_*ex*_*A*_*em*_ (red) and *DA*_*ex*_*A*_*em*_ (purple) streams in the 48-spot setup. The sensitivity of the A-channel causes a shift in the *E*_*PR*_ and *S* peak positions as discussed in the next Section.

### D. E-S histograms

Fig. 6 shows E-S histograms for the dsDNA sample with a 12 bp D-A separation, obtained after the burst search and size selection described in Section III B.

The D-only and FRET populations, top left corner and center respectively in the E-S histogram, are clearly distinguishable in all spots. Moreover, as shown in Fig. 7, the FRET population(s) are easily isolated by applying a second burst selection using a minimum threshold on the *DA*_*ex*_*A*_*em*_ counts. A second example of such a selection is shown in Fig. 11. Separation of FRET species from singly-labeled species is the primary advantage of dual laser excitation^17,39^.

Even without any calibration, the spread across different spots is limited and does not affect the ability to distinguish subpopulations. This is evident in Fig. 8, which shows the *E*_*PR*_ and *S*_*u*_ peak center position in different spots for both D-only and FRET populations. Fig. 9 shows the center and ±1*σ* range from Gaussian fits of *E*_*PR*_ and *S*_*u*_ histograms of the FRET population (blue dot and error-bar). The orange dot is the mean center peak position of the FRET population across all spots. For most spots, the deviation of the peak position is well below the ±1*σ* range, with the exception of spot 19, where the lower A-pixel PDE causes a larger deviation. Note that Fig. 8 and 9 present results without calibration, thus showcasing the minimum performance of the system. It is possible to virtually eliminate spot-to-spot variations of the E-S peak position during post-processing by applying a spot-specific calibration, as briefly described in the next Section (details in Appendix E). As an additional example, the E-S histogram for a low FRET dsDNA (22 bp D-A separation) is reported in Fig. 15, Appendix F.

### E. Pooling data from all spots

The final step of multispot analysis consists in merging data from all spots in order to increase the effective data accumulation rate. Non-uniformities between different spots can be accounted for by applying a two-step correction, in which each correction factor γ and *β* (Eq. D3 and D4) is decomposed into a product of two factors applied successively: an average correction factor computed over all spots and a spot-specific relative correction. The spot-specific correction of γ and *β* can be easily computed from a measurement of a static FRET sample (details in Appendix E). For simplicity and due to the good uniformity among spots, we do not apply any spot-specific corrections in this work.

Fig. 10 shows the cumulated E-S histograms of a mixture of two dsDNA constructs with D-A separation of 12 and 22 bp (see Section III A), resulting in mean *E*_*PR*_ of ~ 0:6 and ~ 0:15 respectively. A significant D-only population is visible as a peak close to *E*_*PR*_ = 0:05, *S*_*u*_ = 1. Due to the relatively low PDE of the acceptor channel (see Section II A), the separation of D-only and FRET populations could be problematic in principle. As previously shown (Fig. 7), FRET populations can be selected by setting a threshold on background-corrected *DA*_*ex*_*A*_*em*_ counts. Fig. 11 shows that this selection effectively removes the large D-only peak, isolating the FRET populations without significant loss of FRET bursts.

**Figure 4.**
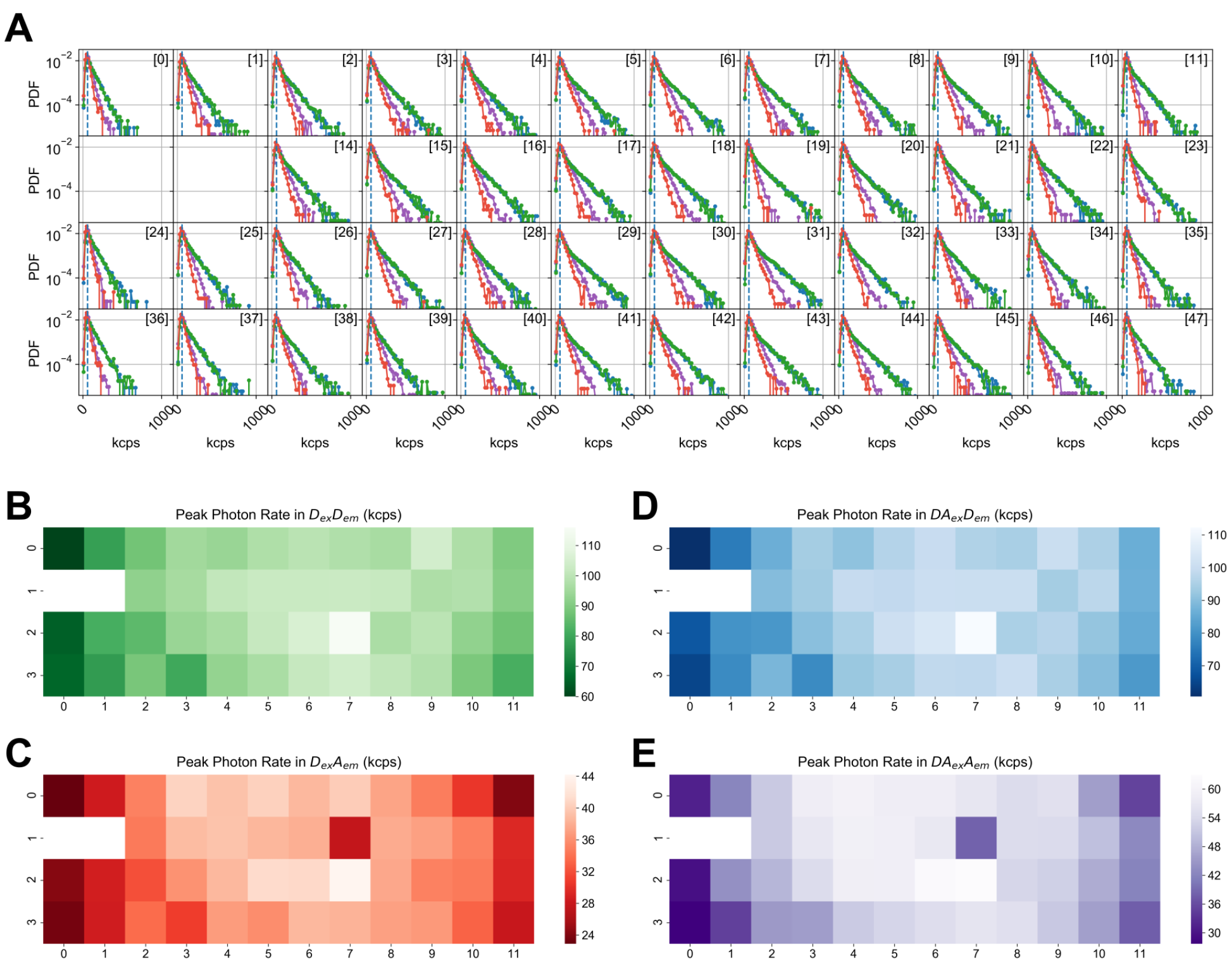
Peak burst photon rates in each of the 48 spots for a dsDNA sample with a 12 bp D-A separation. The output laser powers measured before any optics were set to 200 mW and 400 mW for the D- and A-laser respectively. A: Full distribution of peak photon rates. B-E: Mean of the peak photon rate distribution in different photon streams. Two lateral spots in the second row exhibit no signal because of two malfunctioning pixels in the D-SPAD array. Colors correspond to different photon streams. Green: *D*_*ex*_*D*_*em*_, red: *D*_*ex*_*A*_*em*_, light blue: *DA*_*ex*_*D*_*em*_, purple: *DA*_*ex*_*A*_*em*_. See appendix D for ALEX and PAX streams definitions. For more details see the accompanying 48-spot PAX analysis notebook^37^.

In multispot measurements, it is in principle possible that a molecule detected in one spot could be detected in a different spot, potentially affecting the conclusions drawn from the cumulative data from all spots. Two possible scenarios need to be considered: (i) a single-molecule signal detected in one spot is detected in another due to crosstalk effects, and (ii) a single-molecule detected in one spot diffuses away and is later detected in another spot. The first scenario can be excluded due to the geometry of the setup which ensures that there is no overlap between nearby detection volumes and due to the low optical crosstalk coefficients of the type of SPAD array detectors used in this work^16^. The second scenario would not affect burst analysis as these events would only result in a correlation between bursts in different spots over a timescale determined by diffusion from one spot to the other (Δ*t* ≫ *d*^2^/4*D*, with *d* ≈ 5 μm, *D* ≈ 100 μ*s*^2^*s*^−1^, yields Δ*t* ≫ 60 ms for two nearest-neighbor spots). In practice, we were unable to detect any such cross-correlation in the measurements reported here, suggesting that, due to the size of and the separation between excitation spots, the probability of occurrence of such events is very low.

As an illustration of the high-throughput capabilities of our setup, Fig. 12 and 13 show an example from 5 s-long acquisition windows obtained with the same doubly-labeled dsDNA sample droplet, successively using the single-spot μsALEX and multispot setups (the sample is the 12 bp D-A separation dsDNA described in Section IIIA). In both cases, a constant rate-threshold (20 kcps) burst search was used, followed by burst selection based on *D*_*ex*_ counts (⋀_*γ*_ > 20, see eq. D5). The particular time windows illustrated in Fig. 12 and 13 shows a 37-fold difference in burst numbers between the two measurements (40±11-fold computed over all consecutive 5 s windows within the two 5 min acquisitions), reasonably consistent with the 46-fold difference expected between the two setups, assuming that all experimental conditions are identical. The large standard deviation of the measured ratio reflects the naturally large variance of the burst rate (number of bursts per unit time) within any given measurement. Moreover, since the burst rate depends on a number of measurement and analysis parameters, the burst rate ratio should not be given excessive significance. For instance, the lower power in the lateral spots of the multispot setup (see Fig. 3) will result in smaller bursts and therefore less bursts surviving the selection. While the lower detection efficiency of the SPAD array in the red region of the spectrum (compared to the single spot setup, see Section II A) reduces the acceptor signal in the multispot measurement, also resulting in smaller and fewer bursts above the size selection threshold, this potential source of differences was compensated by the use of *γ*-corrected burst sizes (eq. D5). Finally, differences in observation volumes between setups, as indicated by the slightly longer burst durations in the multispot measurement (suggesting larger volumes), will also translate in different detected burst rates. Despite these caveats, it is obvious that the multispot setup detects a much larger number of bursts than the single spot setup.

**Figure 5.**
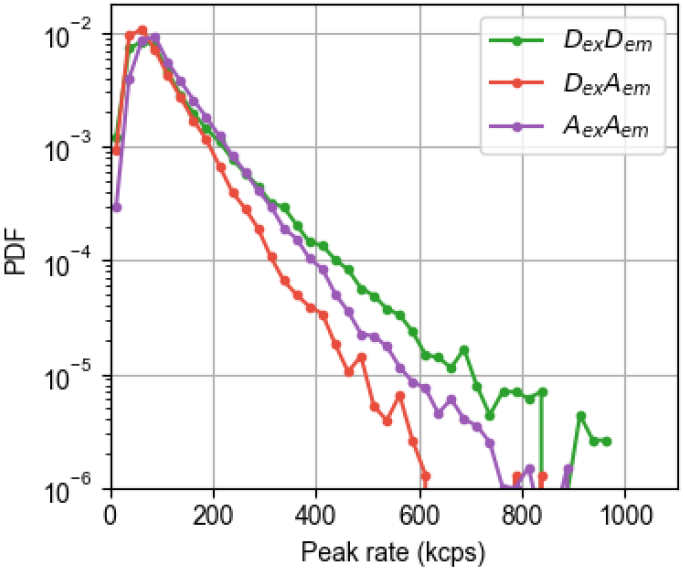
Distribution of peak photon rates in a single-spot μsALEX measurement of the same dsDNA with a 12 bp D-A separation used in 48-spot measurements (Fig. 4). Average laser powers entering the microscope after AOM alternation were 190 μW and 80 μW. Colors correspond to different photon streams. Green: *D*_*ex*_*D*_*em*_, red: *D*_*ex*_*A*_*em*_, purple: *A*_*ex*_*A*_*em*_. See appendix D for ALEX and PAX streams definitions. For more details see the accompanying single-spot ALEX analysis notebook^38^.

Comparison of Fig. 12 and 13 shows that a significant number of bursts (~1,000) can be obtained in only a few seconds of acquisition. Such a high throughput is advantageous for stop-flow or equivalent, real-time kinetic measurements, where a reaction is triggered at time zero and the sample’s evolution monitored continuously afterward. The measurement’s temporal resolution, *i.e.* the smallest usable time window (or time bin), depends inversely on the burst rate (as well as on the type of information to be extracted from the data), based on number statistics. In the simple case where the initial and final FRET states of a reaction are known, and the respective amount of each population is an appropriate reaction parameter, the number of burst in each subpopulation can be extracted from the FRET histogram with a relatively small number of bursts per time bin. The number of bursts collected in the experiment reported in Fig. 12 would definitely be compatible with an effective time resolution ≤ 5 s. To fully take advantage of such a temporal resolution, the experimental dead time (duration of the initial mixing step in the reaction) would need to be shorter, and may require an automated microfluidic system (≪1s, by contrast with the manual mixing reported in the kinetic measurement of ref. 16, where a deadtime of at least 15-20 s was obtained). Such a multispot system could also advantageously be used to perform rapid series of measurements of the same sample in different conditions, or of different samples, for high-throughput screening applications, among other possibilities.

## IV. CONCLUSION

We have described a 48-spot, 2-laser excitation setup designed for high-throughput smFRET assays. Compared to our previous multispot setup^16^, the number of spots was increased six-fold with a corresponding increase in throughput. While larger SPAD arrays have been demonstrated by other groups, they are fabricated using standard high voltage CMOS processes resulting in poorer photon counting performance than the custom technology process employed here. Convincing applications for cell FCS and FLIM, among others, have been published with these CMOS SPAD arrays^42–46^, (for a comprehensive review see 47) but they still remain far from providing the sensitivity needed for single-molecule applications.

Compared to our previous works^16,48^, a second alternating excitation laser was incorporated, and the corresponding alignment hurdles were solved, permitting sorting of single-molecules according to their D-A stoichiometry. In particular, we have shown that the setup allows identifying singly and doubly-labeled species over the full range of FRET efficiencies, opening the door to a much wider range of assays than was previously possible.

We presented a detailed description of the multispot setup and alignment procedure, which incorporates a number of technical solutions of potential interest for other applications. We also illustrated the sm-FRET measurement capabilities of the new setup using doubly-labeled dsDNA molecules as a proof of principle demonstration of sub population separations and high-throughput measurements. Finally, we provided a comparison of its performance with a standard single spot (confocal) μsALEX setup. Applications of this new instrument to the study of the initial stages of bacterial transcription and high-throughput diagnostics will be explored in future work.

**Figure 6.**
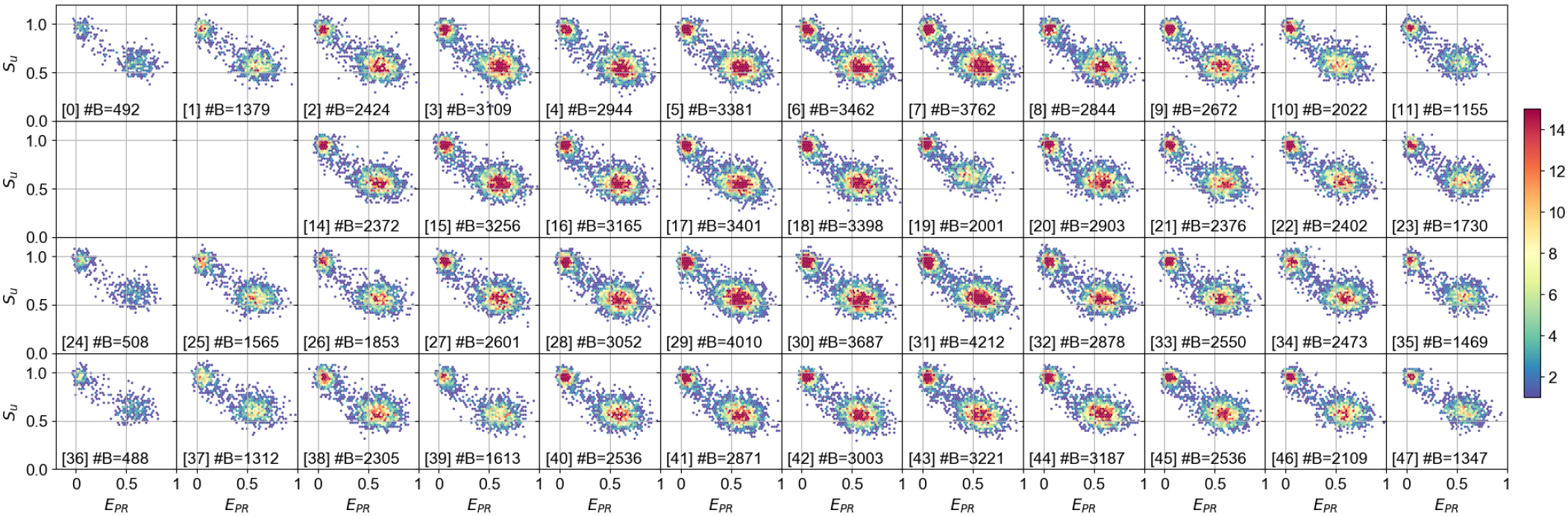
*E*_*PR*_ versus *S*_*u*_ histograms in the different spots for the dsDNA sample with a 12 bp D-A separation. Two subpopulations are visible: D-only (approximately *E*_*PR*_ = 0, *S*_*u*_ = 1) and FRET population (approximately *E*_*PR*_ = 0:6, *S*_*u*_ = 0:6). Burst search was performed using all photons with a constant threshold (50 kcps). Burst selection was performed on the total burst size after background correction, using a threshold of 40 photons. The legend in each subplot reports the spot number in brackets and number of bursts (#B).For more details see the accompanying 48-spot PAX analysis notebook^37^.

**Figure 7.**
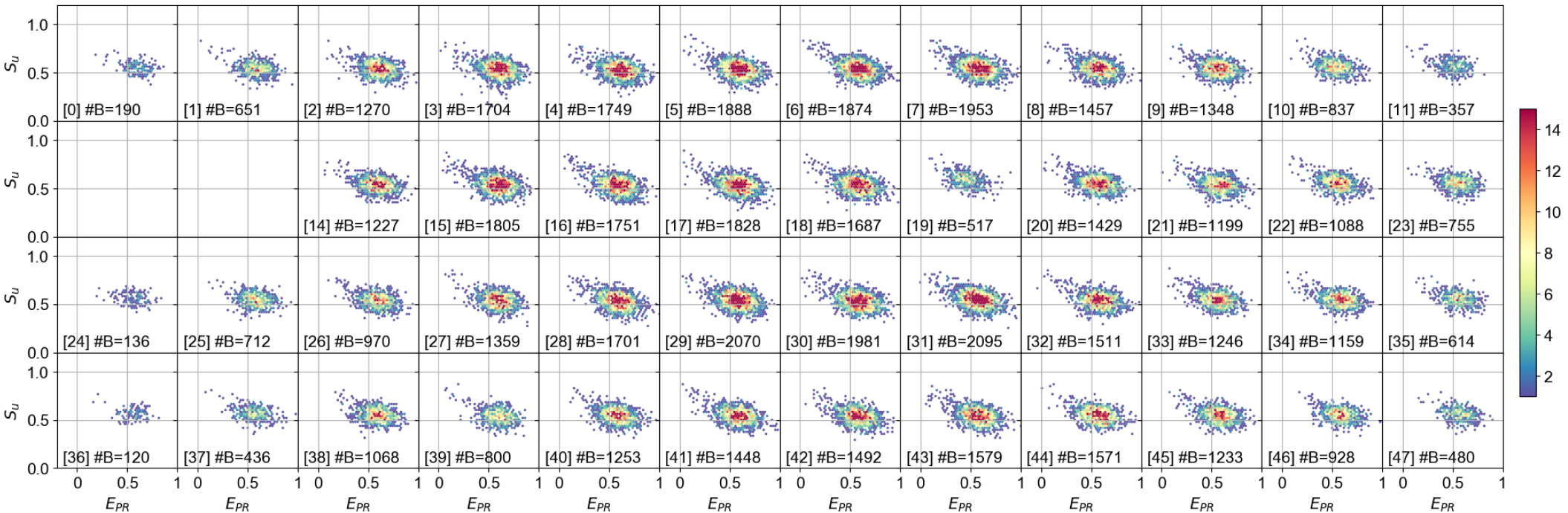
*E*_*PR*_ versus *S*_*u*_ histograms in the different spots for the dsDNA sample with a 12 bp D-A separation. Data analysis and burst search are identical to figure 6, while burst selection is tailored to select only the FRET population: a burst is selected if the number of counts in the *D*_*ex*_*A*_*em*_ and *DA*_*ex*_*A*_*em*_ streams are both larger than 20. The legend in each subplot reports spot number in brackets and number of bursts (#B).For more details see the accompanying 48-spot PAX analysis notebook^37^.

## ACKNOWLEDGMENTS

The authors thank Luca Miari for help in the initial stage of this project, Mr. Yazan Alhadid and Dr. Eitan Lerner for help with single molecule sample preparation and Dr. Eitan Lerner for critical reading of the manuscript. We thank Dr. Bentolila for the generous loan of a LCOS-SLM from the Advanced Light Microscopy/Spectroscopy Shared Resource Facility at the California NanoSystems Institute at UCLA. Research reported in this publication was supported by the National Institute of General Medical Sciences of the National Institutes of Health under Award Number R01 GM095904 & R01 GM069709, and by the National Science Foundation under Award Number MCB 1244175. The content is solely the responsibility of the authors and does not necessarily represent the official views of the National Institutes of Health or the National Science Foundation.

Conflict of interest statements: S. Weiss discloses intellectual property used in the research reported here. M. Ghioni discloses equity in Micro Photon Devices S.r.l. (MPD). No resources or personnel from MPD were involved in this work. The work at UCLA was conducted in Dr. Weiss’s Laboratory.

## Appendix A: Detailed setup description

The setup (Fig. 1) comprises two excitation CW lasers emitting at 532 nm and 628 nm (2RU-VFL-Series, MPB Communications Inc., QC, Canada). For each laser, a half-wave plate and polarizing beam splitter are used for polarization and intensity control, as the polarization orientation must be aligned along the direction required by the LCOS-SLM. The 628 nm laser beam passes through an AOM (P/N 48058 PCAOM, electronics: P/N 64048-80-.1-4CH-5M, Neos Technology, Melbourne, FL) used for μs time-scale modulation. The 532 nm laser is not modulated. Each laser beam goes through a first beam expander (Keplerian telescope, doublet lenses: 50 mm and 250 mm focal lengths). Two periscopes bring the beams to a raised optical breadboard where an inverted microscope body (IX-71, Olympus Corp., Waltham, MA) stands, its bottom port sitting over a circular aperture in the breadboard. Beyond the periscope, each beam goes through a second adjustable beam expander (3X, P/N 59-131, Edmund Optics Inc.). The red laser beam is reflected off mirrors *M*1_*R*_ and *M*2_*R*_ and phase-modulated by the “red” LCOS-SLM (P/N X10468-07, Hamamatsu, Japan), before passing through the dichroic mirror *D*_*MIX*_. The green laser beam is reflected off *M*3, is phase-modulated by the “green” LCOS-SLM (P/N X10468-01, Hamamatsu) and combined with the red excitation via the dichroic mirror *D*_*MIX*_ (T550LPXR, Chroma Technology Corp, VT). Both beams are recollimated by the *L*_3_ lens (f = 250 mm, AC508-250-A, Thorlabs) and focused into the sample by a high-NA water immersion objective lens (UAPOPlan 60X, NA 1.2, Olympus) after being reflected off the excitation dichroic mirror *DM*_*EX*_ (Brightline FF545/650-Di01, Semrock Inc., NY). The excitation pattern forms a dual-color 12x4 array of spots in the sample, matching the geometry of the two SPAD arrays. The fluorescence emission is collected by the same objective lens, passes through the excitation dichroic *DM*_*EX*_ and is focused by the microscope’s tube lens *L*_2_ on either the side or bottom port of the microscope. The side port is equipped with a CMOS camera (Grasshopper3 GS3-U3-23S6M-C, FLIR Integrated Imaging Solutions Inc., BC, Canada) used during alignment, while the bottom port redirects the beams toward the SPAD array emission path. Here, a relay lens *L*_4_ (f = 100 mm, AC254-100-A, Thorlabs) recollimates the image and sends it to an emission dichroic mirror *D*_*EM*_ (Brightline Di02-R635, Semrock), which splits the signal into donor (D) and acceptor (A) spectral bands. The D signal goes through a band-pass filter (Brightline FF01-582/75, Semrock) which removes residual 628 nm laser leakage and helps suppress Raman scattering from the 532 nm laser. Both D and A signals are refocused by lenses *L*5_*D*_ / *L*5_*A*_ (f=150 mm, AC254-150-A, Thorlabs) on two 48-pixel SPAD arrays^29^ (denoted as D and A-SPAD in the text).

**Figure 8.**
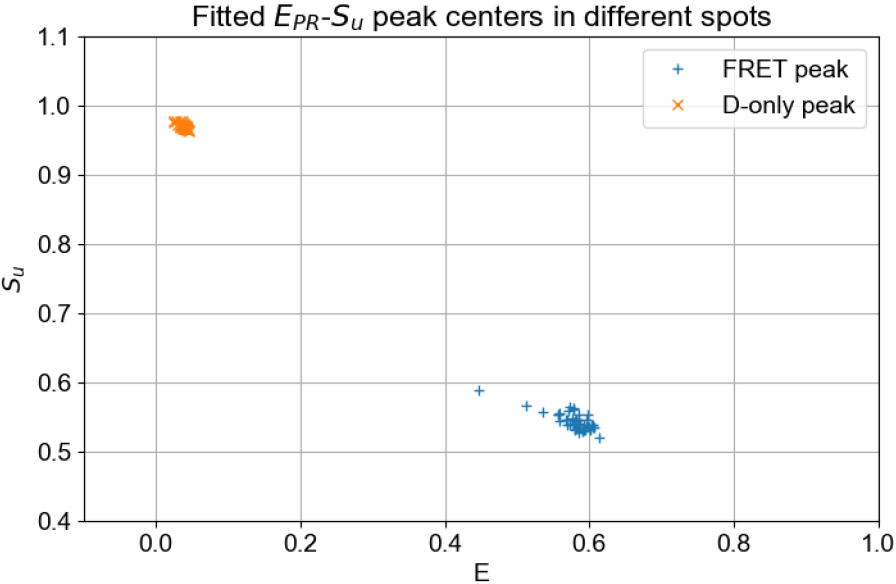
Scatter plot of the fitted *E*_*PR*_, *S*_*u*_ peak position in the different spots for the D-only (*orange cross*) and FRET populations (*blue plus*). Values were obtained by Gaussian fit of the 1-D histogram of *E*_*PR*_ and *S*_*u*_ after a bursts selec-tion that isolated D-only and FRET populations, respectively. For more details see the accompanying 48-spot PAX analysis notebook^37^.

Both SPAD arrays are mounted on 3-axis micropositioners. Motion along the X & Y directions orthogonal to the optical axis are software-controlled via open-loop piezo-actuators (P/N 8302; drivers: P/N 8752 & 8753; Newport Corporation, Irvine, CA). The third axis (Z) uses a manual actuator, as requirements on the Z direction are much less stringent than for the X & Y directions. The D-SPAD array is mounted on an additional rotation stage about the optical axis, which is used to match the relative orientation of the SPAD arrays. Software for controlling the micro-positioners is available in the picomotor repository in 25.

Each SPAD array module is equipped with an internal FPGA (Xilinx Spartan 6, model SLX150), a humidity sensor, and a USB 2.0 connection. The default FPGA firmware used in this work allows acquisition of low-resolution (10-100 ms) time-binned counts via the USB connection, and is also used for humidity monitoring. In addition, a standard SCSI connector includes 48 independent outputs providing a pulse for every detected photon in each pixel^29^. The two SCSI ports are fed through a custom adapter to an FPGA-based acquisition board (FPGA board: PXI-7813R, PXI rack: PXI-1000B, National Instruments, Austin, TX) which performs photon time-stamping with 12.5 ns resolution in parallel on the 96 channels (task implemented in LabVIEW using the LabVIEW FPGA Module, code available in the Multi-channelTimestamper repository in ref. 25). The FPGA board transfers data asynchronously to a host PC via an MXI-4 link to a custom acquisition program written in LabVIEW (PXI rack board: PXI-8331; PC board:PCI-8331, National Instruments). The acquisition program also controls the red laser alternation using a pulse generation board (PXI-6602, National Instruments) with a clock synchronized to the time stamping FPGA board through the PXI rack.

In addition to the aforementioned acquisition program, the host computer runs a second LabVIEW program controlling the phase pattern on the two LCOS-SLMs. During alignment, the acquisition program communicates with the LCOS-control program to scan the positions of the LCOS pattern while recording signal from the SPAD arrays (see Appendix B).

Raw data transferred from the FPGA is saved to disk in a binary file together with a text-based metadata file containing measurement details (sample description, laser powers, alternation info, etc.). Both files are used to create the final Photon-HDF5 file^28,49^. Once the measurement is saved on the host PC, the raw data is automatically transferred to a Linux-based workstation via 1 Gb Ethernet link. The second workstation automatically performs conversion to Photon-HDF5 and data analysis, leaving the host PC available for acquiring the next set of data. The scripts for data transfer, conversion and automated analysis are available in the transfer_convert repository in ref. 25.

**Figure 9.**
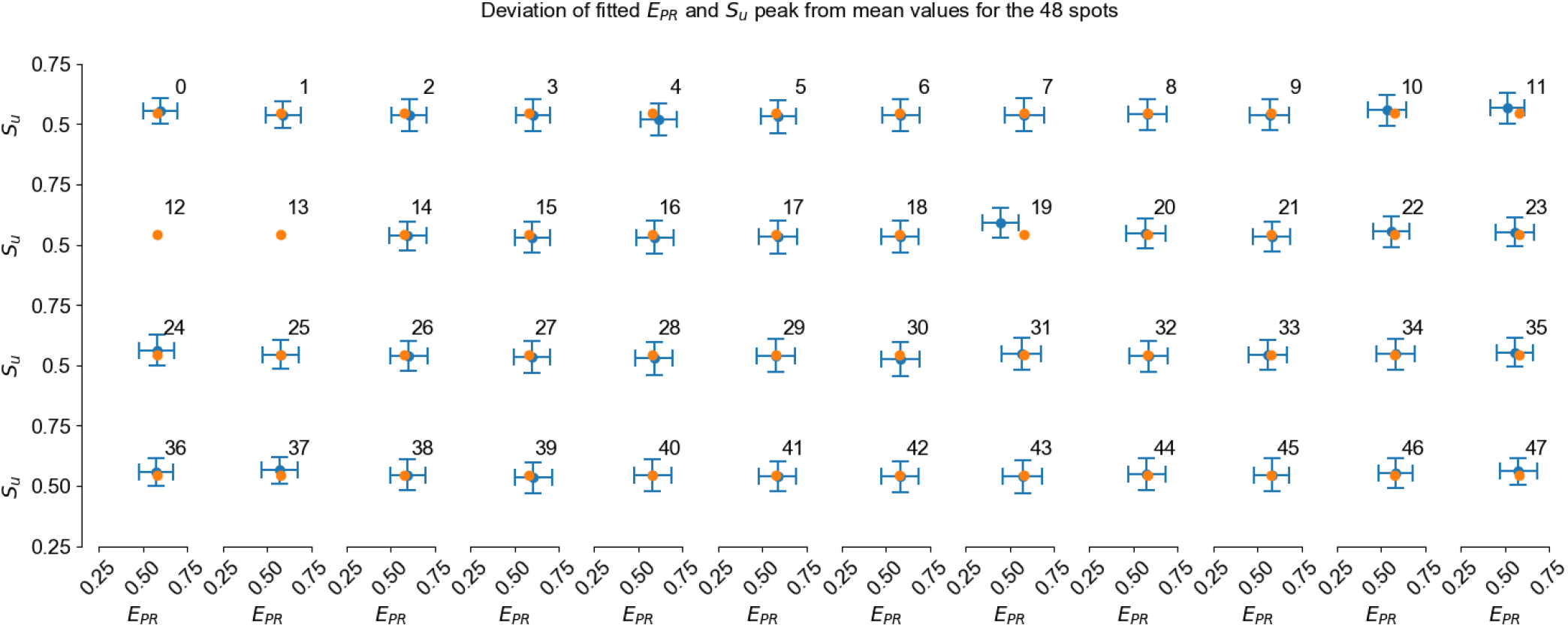
Fitted FRET peak position (*E*_*PR*_, *S*_*u*_, *blue dots*) and ±1*σ* of the fitted Gaussian (*blue error bars*) for the 46 active spots. As a reference, the mean *E*_*PR*_, *S*_*u*_ across all 46 spots (*orange dot*) is reported in each subplot. The spot number is indicated in the toPRight corner of each subplot.For more details see the accompanying 48-spot PAX analysis notebook^37^.

**Figure 10.**
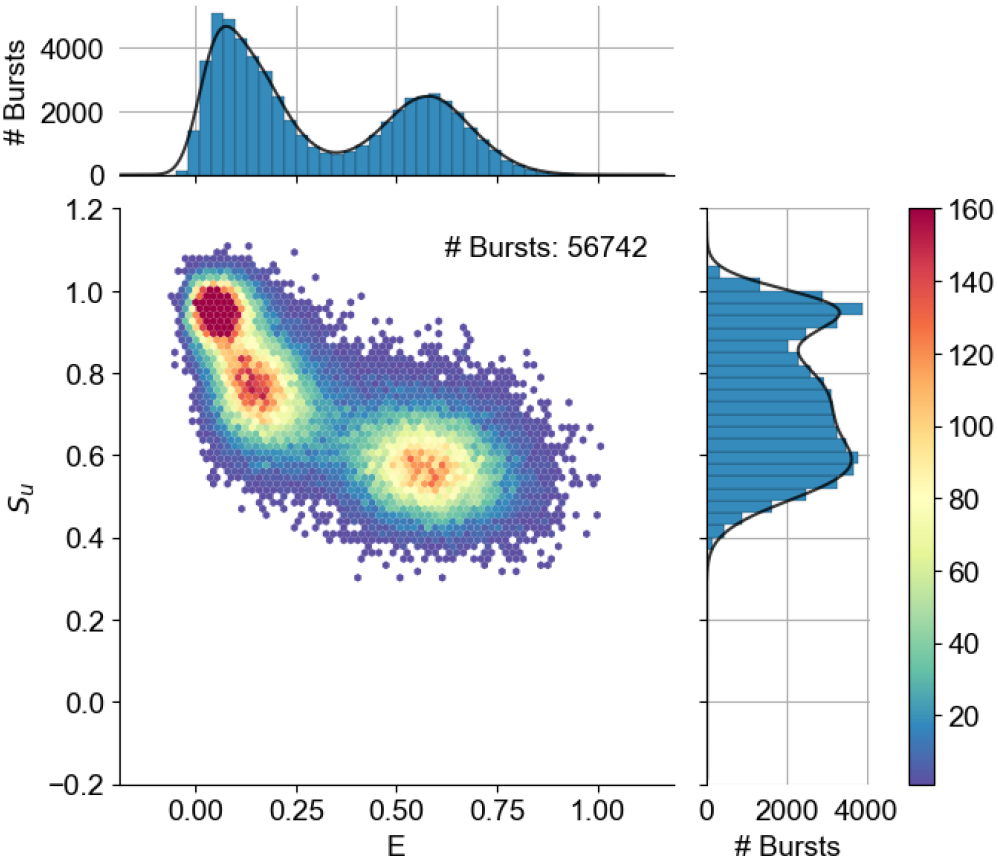
48-spot PAX measurement of a mixture of two dsDNA constructs with 12 and 22 bp D-A separation (sample details in Section III A). Burst search was performed on all channels using a constant-rate threshold of 50 kcps. Burst were selected based on their total size with the criterion ∧_*γP AX*_ > 80, using γ = 0:5 (see eq. D14). The E-S histogram was built by pooling data from all spots, without spot-specific correction. For more details, see the accompanying notebook for the 48-spot PAX analysis of the 12 and 22 bp mixture^40^.

## 1. LCOS-SLM Modulation

The array of 48 excitation spots is generated separately for each color by two LCOS-SLMs via phase modulation of an incoming plane wave, as previously described in 16 and 32. Briefly, the LCOS-SLM implements the phase profile of a lenslet array which focuses the incoming plane wave into an array of spots 3-4 cm in front of the LCOS-SLM surface (see Fig. 1). A rectangular region of the LCOS-SLM is subdivided into 12x4 adjacent blocks each implementing a single lens. The pattern can be adjusted by changing its center position, rotation, and X and Y pitch independently (operations equivalent to shifting, rotating or scaling the lenslet array). For both excitation wavelengths, the pitch and therefore the diameter of the lenslets is imposed by the detector geometry and the magnification of the optical setup in both excitation (83×) and emission (90× = 60 × 1.5) paths. Nominally, the spot pitch in the sample matching the detector pitch is 5.5 μm(500 *μ*m/90) in both direction, resulting in an LCOS-SLM lenslet pitch of 463 μm (23.1 LCOS-SLM pixels). The value is optimized during alignment to match the actual magnification and optical aberrations. Keeping constant the LCOS-SLM lenslet diameter and pitch, a change in the lenslet focal length results in a change in NA and therefore spot size. The ratio of focal lengths in the two LCOS-SLM (32 mm for the red and 36 mm for the green) is chosen to compensate the difference in PSF sizes between 532 nm and 628 nm wavelengths. Note that changing the lenslet focal length requires changing the distance between *L*_3_ and the LCOS-SLM so that the LCOS focal plane remains at focal distance from *L*_3_.

**Figure 11.**
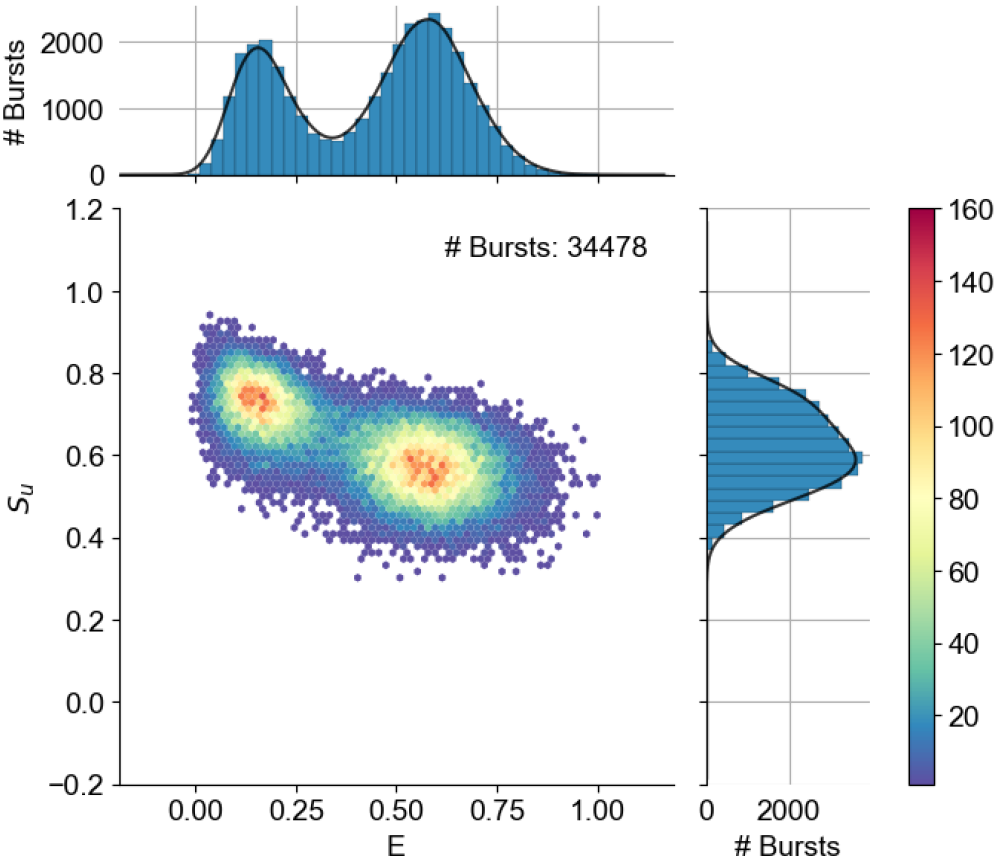
48-spot PAX E-S histograms of the same measurement of Fig. 10. Additional filtering of the D-only population was performed using the criterion 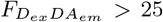 (see Appendix D). The E-S histogram was built by pooling data from all spots without spot-specific correction. For more details, see the accompanying notebook for the 48-spot PAX analysis of the 12 and 22 bp mixture^40^.

**Figure 12.**
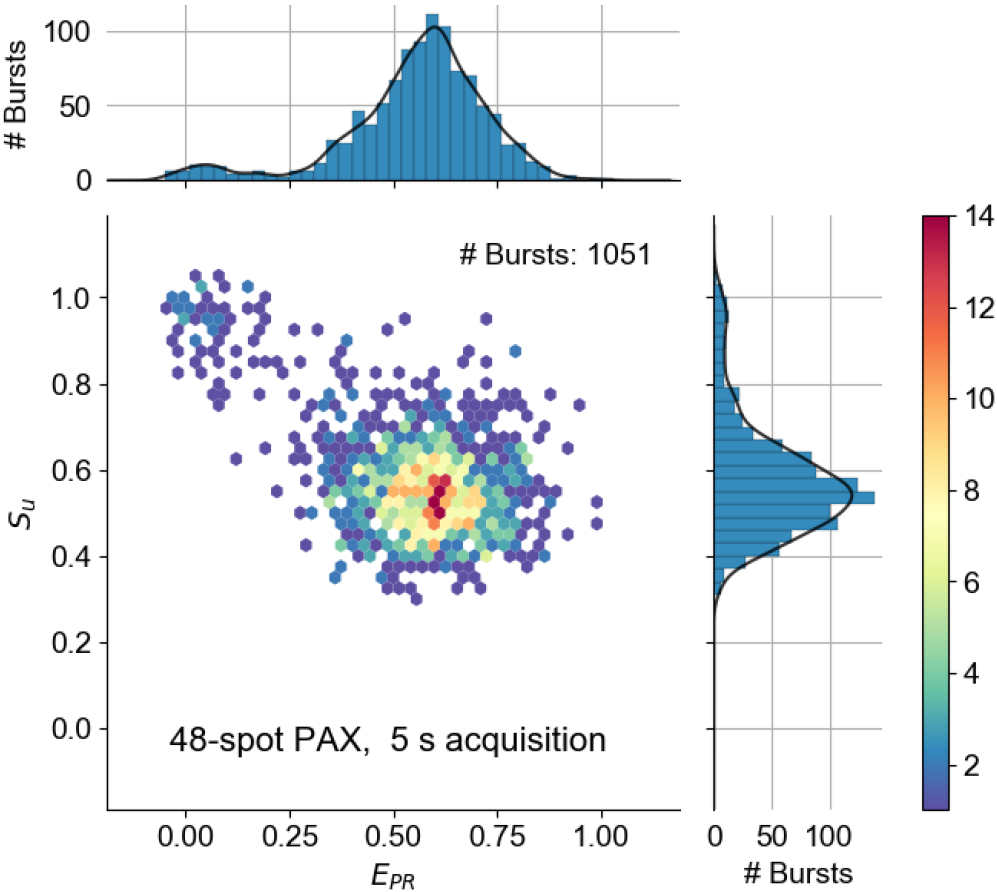
Multispot E-S histogram obtained from 5 s of acquisition by pooling bursts from the 46 active spots. For more details see the accompanying notebook for comparison of a single spot to 48-spots^41^.

**Figure 13.**
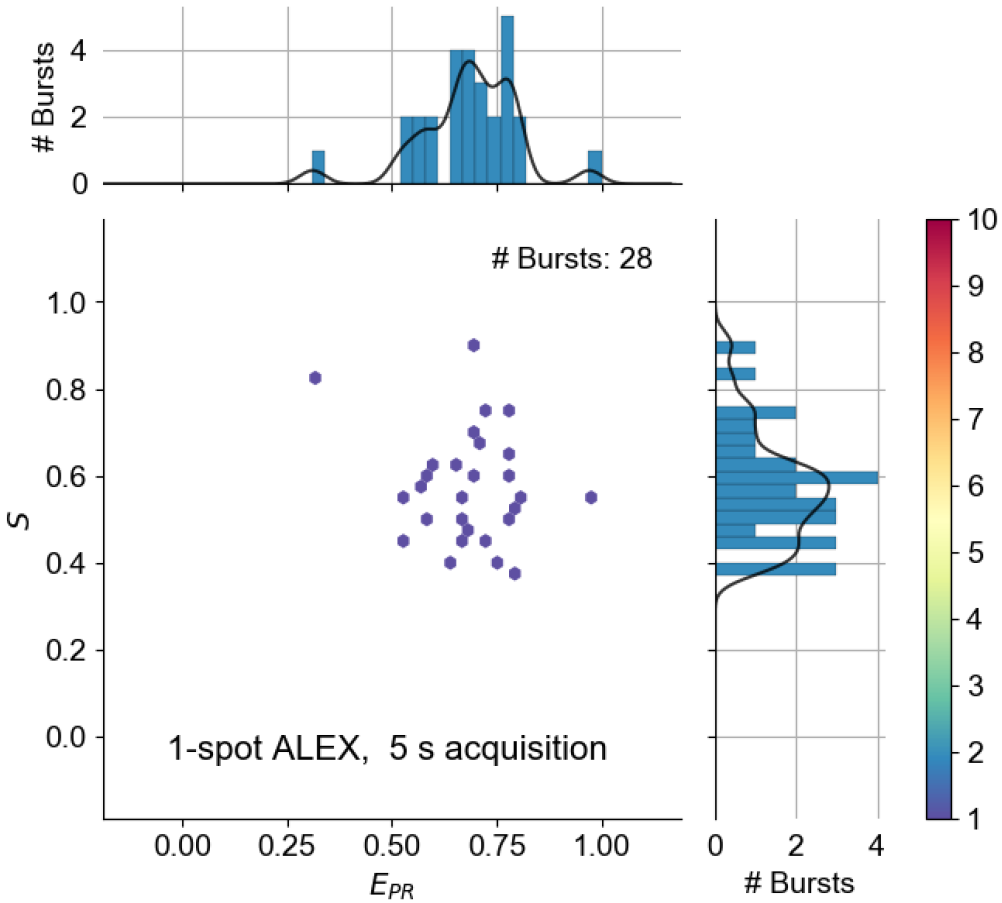
Single-spot E-S histogram obtained from 5 s of acquisition for the same sample as in Fig. 12. For more details see the accompanying notebook for comparison of a single spot to 48-spots^41^.

The LCOS-SLM region surrounding the 12x4 pattern receives light that can can result in stray ":wide-field" excitation and therefore increase the background signal. For this reason, we fill the unused LCOS-SLM area with a "beam steering" pattern (a periodic pattern in one direction) that diffracts the incoming light at an angle with respect to the optical axis. This "steering" ensures that light not contributing to the multispot pattern is not collected by the back aperture of the objective lens. Additionally, the expanded laser beam is clipped by two rectangular apertures (slits) approximately 1 mm larger than the multispot pattern, further reducing sources of background. This approach achieves low background without the need of an additional spatial filter as was used in our previous 8-spot setup^16^.

A similar approach for multispot generation was used for multi-confocal FCS by Kloster *et al.*^33^. The fundamental difference of their method is the use of a much longer LCOS focal length to construct a single phase pattern for all spots (as the sum of the contributions of each single spot). By contrast, in our approach, different portions of the LCOS-SLM are allocated to different spots. A detailed experimental comparison highlighting the relative strengths of these two approaches is currently lacking.

Software to generate the multispot phase pattern used in this work is available in the lcos_multispot_pattern repository in ref. 25.

## Appendix B: Laser alignment

Each of the two lasers needs to be aligned in order to ensure (a) maximum uniformity between spot intensities (b) minimal aberrations across the pattern. To achieve (a), the Gaussian laser beam is expanded so that only the central part of the beam covers the excitation pattern (which has a maximum extension of 5 mm). To ensure (b), the geometrical center of the pattern needs to be placed on the optical axis.

In addition, (c) the excitation pattern of the two lasers must be aligned such that there is a maximum overlap between D and A excitation volumes for each spot.

## 1. Individual laser alignment

The 3X beam expanders have an adjustment ring used to control beam collimation. A simple way to ensure beam collimation is by sending the beam into the microscope through the excitation dichroic mirror, removing the external recollimation lens *L*_3_ and the objective lens, while placing a mirror on the sample holder and using the LCSO-SLM as a mirror, *i.e.* displaying a constant phase pattern. Using the camera on the microscope output port, we adjust the collimation until a tight spot is formed. After adjusting the collimation, each beam must be aligned so that the peak intensity is at the center of the optical axis. To this end, after removing the recollimating lens *L*_3_, an iris *I*_2_ is placed before the beam enters the microscope side port. Using an aperture of 1-2 mm, only a narrow beamlet goes through the objective and generates a spot from the cover-glass reflection. Only when the input beam is parallel to the optical axis, the spot will be located in the center of the cross-hair in the microscope’s eyepiece. In order to make the input beams parallel to the optical axis, the last mirrors before the microscope are adjusted (*M*2_*R*_ for the red and *DM*_*MIX*_ for the green laser). When changing the microscope’s focus, we obtain symmetrically concentric patterns only if the input beamlet intersects with the optical axis at the back aperture of the objective lens. Since the direction is already fixed, we move the *I*_2_ iris to obtain the most radially-symmetric defocused pattern. In this way, the beamlet that goes through *I*_2_ coincides with the microscope’s optical axis. The last step involves translating the input beam without changing its incidence angle until the intensity peak is aligned to the iris center. A pure translation is achieved by rotating two mirrors in opposite directions so that the initial and final beam angle remains unchanged. Alignment of beam direction and iris must be repeated until convergence. Once complete, both beams are parallel and concentric with the optical axis to a good approximation. When placing *L*_3_ a spot is formed at a different focus position. *L*_3_ can be aligned by ensuring that this spot is located at the same position as the spot obtained without *L*_3_.

## 2. Achieving overlap of the green and red patterns

Starting with the green LCOS-SLM, we project a multispot pattern into a highly concentrated solution of Cy3B and ATTO647N dyes (100 nM - 1 μM). Using a square grid with an odd number of spots per side (e.g. 9x9) ensures that one spot is always at the center of the pattern. The camera on the side-port detects an image of the pattern. The centering of the pattern with respect to the optical axis can be assessed from the degree of geometrical aberrations in the lateral spots. We center the excitation pattern by rigidly translating the pattern on the LCOS-SLM so that geometrical aberrations are roughly equivalent on all four sides. Next, we perform a 2D Gaussian fitting of each spot, and from the distribution of waist size and tilt angle for each Gaussian, we estimate a more accurate position of the optical axis (for the analysis see the LCOS pattern fitting notebook^34^). This step may be repeated multiple times until convergence. From this point on, the X & Y positions of the green LCOS-SLM is not changed anymore, and its center becomes a reference for the optical axis position.

Next, we activate the red LCOS-SLM and project a multispot pattern excited by the 628 nm laser. Using the camera, we align the red pattern to the green one used as a reference. An initial coarse adjustment of the red LCOS-SLM pattern is performed manually by observing the emission pattern on the live camera display. Then, the center position of the red LCOS-SLM pattern is finely adjusted by fitting the spot positions in the green and red images (Fig. 3), taken separately (for implementation details see the LCOS pattern fitting notebook^34^).

Finally, in order to reduce the background due to unmodulated light, two custom-made rectangular slits (aluminum with black finish) are added in the path before each LCOS (*S*_*R*_ and *S*_*G*_ in Fig. 1). The slits are aligned to illuminate only the 12×4 pattern (±1 mm) on the LCOS-SLM (see Appendix A1).

## Appendix C: SPAD arrays alignment

Both detectors must be aligned so that each pixel is optically conjugated to the corresponding excitation volume, i.e. the excitation PSF. The goal is to have pairs of corresponding pixels on the two arrays detecting photons from the same sample volume, *i.e.* the detection PSF. At the same time, in order to maximize signal, the detection PSF must be concentric with the excitation PSF. Achieving this with a 2D arrangement of spots and pixels requires not only aligning the X & Y position of the detectors, as in single-spot measurements, but also aligning the relative rotation of the two SPADs and adjusting the pitch and rotation of the excitation pattern to optimally match the detectors’ geometry.

For alignment, we use a high concentration of a dye mixture (ATTO550, ATTO647N, ~500 nM) excited by both lasers. With such a sample, the 532 nm laser generates fluorescence signal in both D and A channels, while the 628 nm laser only generates a signal in the A channel. At this point, the position of both 532 nm and 628 nm excitation patterns on the LCOS-SLM has already been fixed in order to minimize geometrical aberrations as described in Appendix B. Therefore, the excitation pattern position is used as the reference for aligning the SPAD arrays. Tyndall et al.^50^ have presented an automatic procedure to align a LCOS-SLM multispot pattern to the detector. Here we align the SPADs to the LCOS-SLM pattern.

Starting with the green laser only, both SPADs are manually positioned in X and Y to match the center of the excitation pattern. This is achieved by finding the location of the maximum recorded SPAD counts while moving the detectors.

Next, we perform a more automated procedure for fine alignment referred to as “multispot scan”. A multispot scan involves rigidly translating the multispot pattern on an LCOS-SLM (typically a 4x4 spot pattern) in discrete steps along two orthogonal paths, forming a cross. At the same time, counts from a SPAD array are integrated for each pattern position over 300 ms after each step. During a scan, each emission spot draws a cross path approximately centered on a SPAD pixel. A typical scan covers a range of 10 LCOS-SLM pixels with a step size of 0.4 pixel and is performed sequentially in both X and Y directions. The counts acquired as a function of the LCOS-SLM position and form a peak profile, which is used to estimate the SPAD pixel center positions in LCOS-SLM coordinates. Averaging the SPAD pixel positions, we obtain an accurate estimation for the center of the SPAD array. Ultimately, this procedure yields the offset of each SPAD array with respect to the ideal excitation pattern center. With this information, we move the SPAD arrays to the ideal (X, Y) position using software-controlled piezo-driven micro-positioners. The sequence of multispot scan and SPAD array translation is repeated until convergence. Initially, the two SPAD arrays are aligned with respect to the green LCOS-SLM pattern (532 nm). Next, the position of the red LCOS-SLM pattern (628 nm) is fine tuned to match the position of the A-SPAD array (the D-SPAD array does not detect any signal with 628 nm excitation). The optimal position of the red excitation pattern is determined from a multispot scan performed with the red LCOS-SLM, while counts are acquired with the A-SPAD array, as previously described. After this last step, both red an green excitation patterns, as well as D- and A-SPAD array positions are fixed, completing the setup alignment.

The whole fine alignment procedure is routinely performed at the beginning of each day of measurements and lasts about 30 minutes. Fig. 14 shows the fitted coordinates after fine alignment of the central 4×4 set of pixels in the D and A-SPAD arrays.

## 1. Rotation and pitch adjustment

In the previous Section, we outlined the general fine alignment procedure repeated daily when using the multispot setup. However, when building the setup, additional steps are necessary to (a) align the relative rotation of the two SPAD arrays, (b) determine the best pitch in X and Y for the green and red excitation patterns, and (c) optimize the SPAD position along the optical axis (Z).

To extract rotation and pitch information, we perform a multispot scan followed by an additional analysis step. Specifically, the set of (X, Y) positions of each SPAD pixel obtained from the scan is fitted to a rectangular grid. The fitted grid parameters are: center position, X pitch, Y pitch, and rotation angle. Each SPAD array will generally have a different set of fitted parameters.

To adjust the rotation angle, one of the SPAD arrays (D) is rotated about the optical axis in order to match the angle of the second SPAD, where the rotation angle of each SPAD is obtained from the scan fits. Once the orientations of two SPAD arrays match each other, the rotation stage is locked, ensuring long-term stability of the rotational angle.

To adjust the pitch, information from the scan fits is used to finely tune the X and Y pitch of the LCOS-SLM pattern in order to optimally match both SPAD arrays. Residual X and Y pitch difference of 1-2% are observed due to non-idealities, *i.e.* stigmatisms, in the optical path.

## Appendix D: ALEX and PAX

In ALEX, two alternation time windows, *D*_*ex*_ and *A*_*ex*_ (respectively the D or A excitation window) and two detectors (D and A) are involved. This results in four photon streams noted *D*_*ex*_*D*_*em*_, *D*_*ex*_*A*_*em*_, *A*_*ex*_*D*_*em*_, *A*_*ex*_*A*_*em*_ where the first letter indicates the excitation period and the second the detection channel. The *A*_*ex*_ *D*_*em*_ stream only contains background because there is no fluorescent emission in the D-spectral band during A-laser excitation and is therefore ignored. For simplicity, we assume in the following that all quantities have been corrected for background^36^.

**Figure 14.**
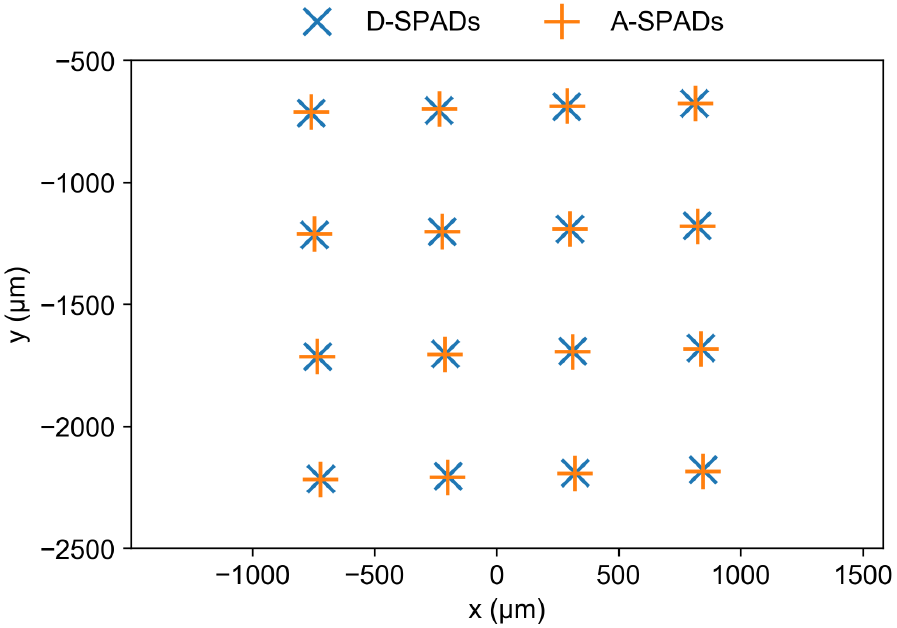
Experimental SPAD pixel coordinates after fine alignment for the D-SPAD and A-SPAD arrays obtained with scans of the green LCOS-SLM. D-SPADs center positions are denoted by ‘X’ and A-SPADs center positions are denoted by ‘+’. The mean distance between D- and A-SPAD pixels is 2.3 μm. For details see the accompanying SPAD alignment notebook^34^.

A PAX setup has two detectors (D and A) but only one alternating laser (A). As in ALEX, two alternation time windows can be defined: *D*_*ex*_, corresponding to the interval during which only the D-laser is on and *DA*_*ex*_, when both lasers are on. As before this results into four photon streams noted *D*_*ex*_*D*_*em*_, *D*_*ex*_*A*_*em*_, *DA*_*ex*_*D*_*em*_, *DA*_*ex*_*A*_*em*_. Formally, the only difference with ALEX is that *A*_*ex*_ in ALEX is replaced with *DA*_*ex*_. In PAX, however, all four photon streams contain fluorescent signal. In particular, *DA*_*ex*_*D*_*em*_ contains D-fluorescence due to D-laser excitation (the corresponding term in ALEX, *A*_*ex*_*D*_*em*_, contains only background). With this notation, we can define the total fluorescence signal during D-excitation (valid in both ALEX and PAX) as:

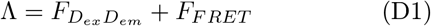

where the *F* quantities are background-corrected photon counts. *F*_*F RET*_ is the detected acceptor fluorescence due to FRET, computed by subtracting from 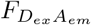 the D-leakage in the acceptor channels (*Lk*) and the A-direct-excitation by A-laser (*Dir*)^24^:

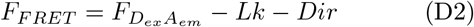

We also need the usual correction factors and *γ* and *β*^24^:

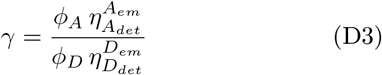

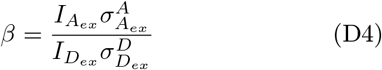

where *ϕ*_*A*_, *ϕ*_*D*_ are the acceptor and donor quantum yields and 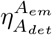, 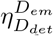 are the detection efficiencies of the D and A signals in the D and A channels. In eq. D4, 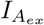 and 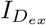 are A and D-excitation intensities, while 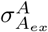 and 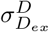 are the dye absorption cross-sections at their respective laser wavelengths. *β* accounts for the difference in D and A-dye excitation rates when each dye is excited by its respective laser.

We can define the *γ*-corrected total signal upon D-excitation as^24,51^:

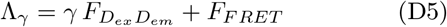

With these definitions, the proximity ratio *E*_*PR*_ and FRET efficiency *E* are given by the following expression, valid for both ALEX and PAX:

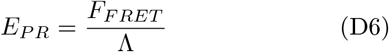

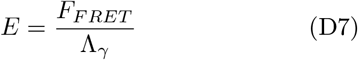

By contrast, the expression for the stoichiometry ratio *S* is slightly different for ALEX and PAX. In ALEX, *S* and its corrected version *S*_*γβ*_ are defined as:

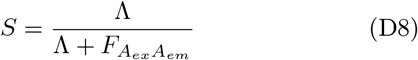

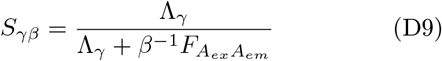

The value *S*_*γβ*_ is always centered around 0.5 for doubly-labeled species, regardless of FRET efficiency or D and A-excitation intensities.

In PAX, there is no such signal as 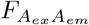. However, when the alternation period is sufficiently short to assume that the excitation intensity doesn’t change from one interval to the next (which is usually the case), an equivalent quantity can be computed by subtracting the contribution of D-excitation to 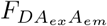:

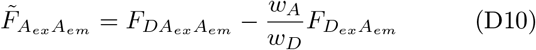

where *w*_*A*_ and *w*_*D*_ are the durations of the *DA*_*ex*_ and *D*_*ex*_ excitation periods respectively. Typically the alternation periods have the same duration (*i.e.* duty cycle = 0.5) and *w*_*A*_ / *w*_*D*_ = 1. Expressions for *S* and *S*_*γβ*_ defined for ALEX (eq. D8 and D9) can then be used for PAX, with the replacement of 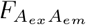 by 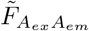:

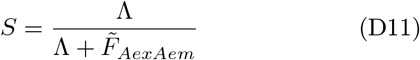

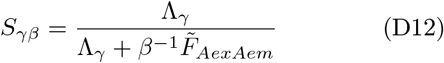

Since in PAX experiments the D-excitation is always on, we can use the signal in 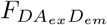 to improve photon statistics. To derive a modified set of PAX expressions for *E* and *S* we start by defining a “modified” total FRET signal as:

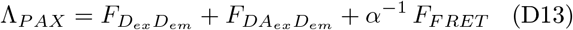

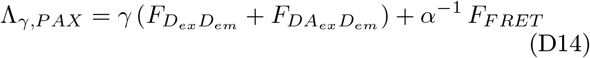

where 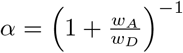 is the *D*_*ex*_ duty cycle (typically, *w*_*A*_ = *w*_*D*_ and *α* = 0.5). The factor *α*^-1^ amplifies the *F*_*F RET*_ signal in order to compensate for the additional donor signal 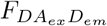.

Based on eq. D13 and D14, we can write modified PAX expressions for *E* and *S*:

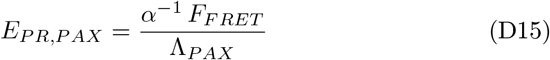

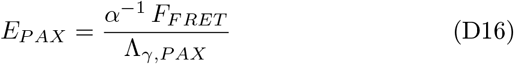

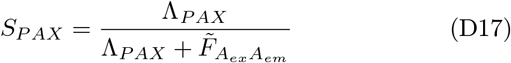

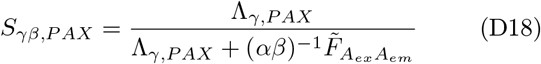

Eq. D15, D16, D17 and D18 contain more photons than the classical expressions and, therefore, can result in lower shot-noise. However, this effect is mitigated by the fact that *F*_*F RET*_ is multiplied by *α*^-1^ to compensate for the additional D signal 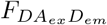, and therefore its shot-noise is amplified.

## 1. Modified stoichiometry

By replacing 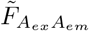 with 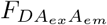 in eq. D11, a modified or “uncorrected stoichiometry” *S*_*u*_ can be defined:

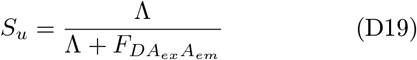

This expression avoids subtracting 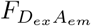 counts from 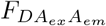 (an operation which sums the statistical noise of these two quantities), therefore improving the overall dispersion of the ratiometric quantity *S*_*u*_. As a result, the Su distributions are narrower, permitting easier separation of FRET and D-only population. Note, however, that Su has a built-in dependency on the population FRET value, in particular *S*_*u*_ decreases with increasing *E*. In this work, even at low FRET values, better separation between FRET and D-only population was achieved using *S*_*u*_ instead of *S*. In general, the advantage of *S*_*u*_ over *S* may change in other situations, especially when signal-to-noise and signal-to-background ratios are large. Once populations are separated in the E — *S*_*u*_ histogram, one can use the classical *S* expression (eq. D11) to compute gamma factor as described in ref. 24. In this work, no attempt was made to recover exact FRET values and D-A distances, therefore no gamma factor calibration was performed. However, we address the issue of differences in collection and detection efficiencies across spots, which can affect such a calibration, in Section E.

## Appendix E: Individual spot corrections

## 1. Gamma correction

The gamma-factor of each spot, *γ*_*sp*_, can be expressed as the product of an average factor *γ*_*m*_ and a spot-specific adjustment factor *χ*_*sp*_:

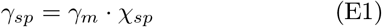

*χ*_*sp*_ can be easily computed from measurable quantities according to the following expression:

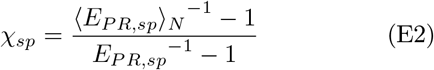

In eq. E2, *E*_*PR,sp*_ is the sub-population proximity ratio measured in a specific spot, and ⟨*E*_*PR,sp*_⟩_*N*_ is its average over all *N* spots (here, *N* = 48).

Eq. E2 follows from the following relation between *E* and *E*_*PR*_^24,51^:

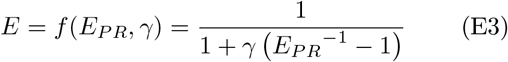

Solving eq. E3 for *γ*, we obtain:

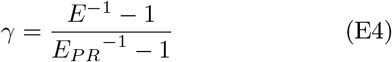

Formally, we can write *γ* = *γ*_1_ *γ*_2_, where *γ*_1_ is associated with a partially corrected proximity ratio *E*_1_ as follows:

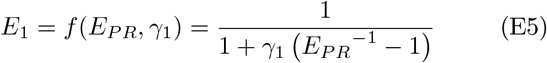

Writing *γ*_1_ as a function of *E*_1_ as in eq. E4 and substituting the expression into eq. E3, we obtain *E* as a function of *E*_1_:

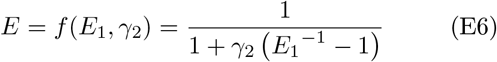

Eq. E6 has the same form as E3 and E6. Therefore, *E* can be obtained by two subsequent (chained) corrections for *γ*_1_ and *γ*_2_ respectively as in eq. E7.

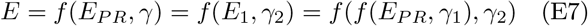

In the multispot case, we apply this property to decompose the gamma correction into a spot-specific correction and an average correction as in E1. In particular, eq. E2 directly derives from E4 with simple substitutions.

## 2. Beta correction

Since, formally eq. D8 and D9 have the same form as *E*_*PR*_ and *E*, we can write an expression equivalent to E3 for *S* and *S*_*γβ*_. Dropping the *γ* subscript, we obtain:

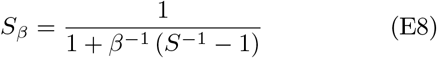

Following the same arguments as in the previous Section, the beta correction can be expressed as the product of a spot-average *β*_*m*_ and an individual spot correction *β*_*sp*_:

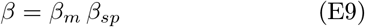

Similarly to eq. E2, we can compute *β*_*sp*_ as:

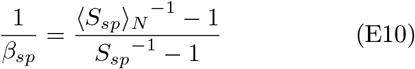

where *S*_*sp*_ is the sub-population non-beta-corrected stoichiometry ratio for a specific spot, and ⟨*S*_*sp*_⟩_*N*_ is the average over all *N* channels (here *N* = 48).

## Appendix F: Additional data

Fig. 15 shows *E*_*PR*_-*S*_*u*_ histograms for the different channels obtained during the measurement of a 22d DNA sample (low-FRET). Due to the choice of donor and acceptor excitation powers during this measurement, the FRET population has a *S*_*u*_ value > 0.5, artificially compressing the histograms in the upper part of the graph. Nonetheless, it is still possible to distinguish FRET from D-only bursts, despite the low value of *E*_*PR*_ for that sample, allowing an unbiased estimation of *E*_*PR*_, as opposed to what would have happened in the absence of acceptor excitation^16^.

**Figure 15.**
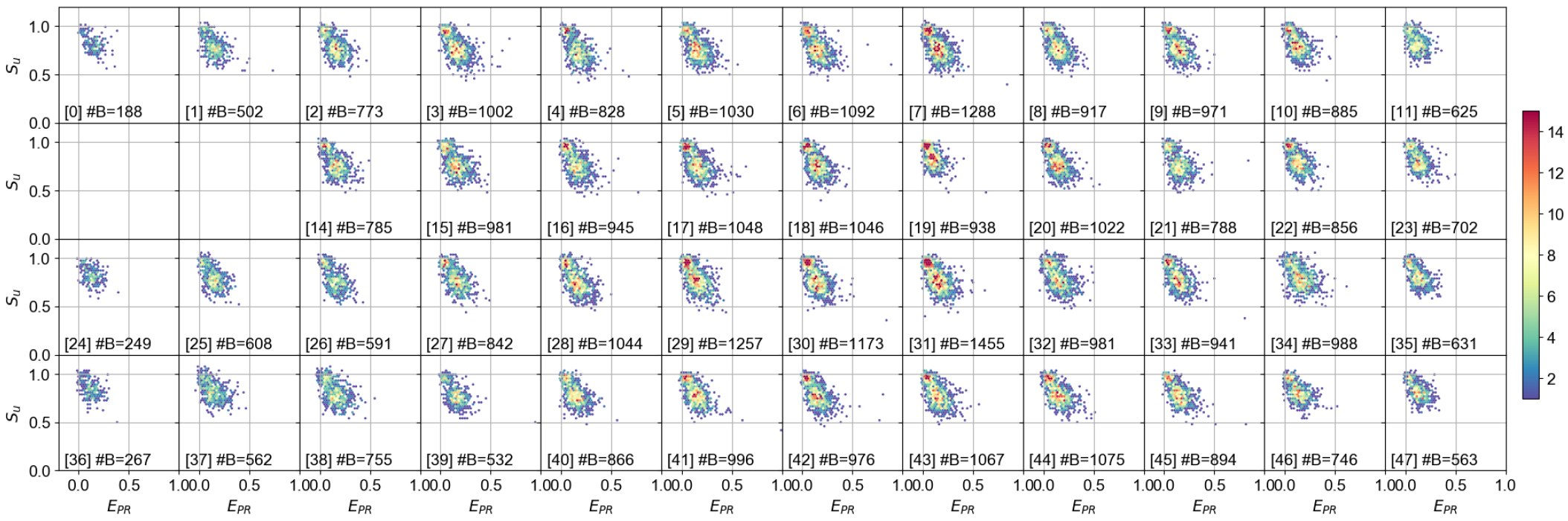
*E*_*PR*_ versus *S*_*u*_ histograms of all spots for the 22d dsDNA sample. Data analysis and burst search are identical as in figure 6. Burst search was performed using all photons with constant threshold (50 kcps). Burst selection was performed on the total burst size after background correction, using a threshold of 40 photons. The legend in each subplot reports spot number in brackets and number of bursts (⧣B). For more details see the accompanying 48-spot PAX analysis notebook^52^.

## REFERENCES

1 M. Dahan, A. A. Deniz, T. Ha, D. S. Chemla, P. G. Schultz, and S. Weiss, Chemical Physics 247, 85 (1999).

2 A. Chakraborty, D. Wang, Y. W. Ebright, Y. Korlann, M. DahanE. Ko-rtkhonjia, T. Kim, S. Chowdhury, S. Wigneshweraraj, H. Irschik, R. Jansen, B. T. Nixon, J. Knight, S. Weiss, and R. H. Ebright, Science 337, 591 (2012).

3 A. N. Kapanidis, E. Margeat, S. O. Ho, E. Kortkhonjia, S. Weiss, and R. H. Ebright, Science 314, 1144 (2006).

4 A. N. Kapanidis, E. Margeat, T. A. Laurence, S. Doose, S. O. Ho, J. Mukhopadhyay, E. Kortkhonjia, V. Mekler, R. H. Ebright, and S. Weiss, Molecular Cell 20, 347 (2005).

5 T. Ha, A. Y. Ting, J. Liang, W. B. Caldwell, A. A. Deniz, D. S. Chemla, P. G. Schultz, and S. Weiss, Proceedings of the National Academy of Sciences 96, 893 (1999).

6 A. A. Deniz, M. Dahan, J. R. Grunwell, T. Ha, A. E. Faulhaber, D. S. Chemla, S. Weiss, and P. G. Schultz, Proceedings of the National Academy of Sciences 96, 3670 (1999).

7 M. Dimura, T. O. Peulen, C. A. Hanke, A. Prakash, H. Gohlke, and C. A. Seidel, Current Opinion in Structural Biology Carbo-hydrate–protein interactions and glycosylation Biophysical and molecular biological methods, 40, 163 (2016).

8 B. Hellenkamp, P. Wortmann, F. Kandzia, M. Zacharias, and T. Hugel, Nature Methods 14, 174 (2017).

9 M. Hoeffling, N. Lima, D. Haenni, C. A. M. Seidel, B. Schuler, and H. Grubmüller, PLOS ONE 6, e19791 (2011).

10 S. Kalinin, T. Peulen, S. Sindbert, P. J. Rothwell, S. Berger, T. Restle, R. S. Goody, H. Gohlke, and C. A. M. Seidel, Nature Methods 9, 1218 (2012).

11 A. Muschielok, J. Andrecka, A. Jawhari, F. Brückner, P. Cramer, and J. Michaelis, Nature Methods 5, 965 (2008).

12 A. Muschielok and J. Michaelis, The Journal of Physical Chemistry B 115, 11927 (2011).

13 J. R. Fries, L. Brand, C. Eggeling, M. Köllner, and C. A. M. Seidel, The Journal of Physical Chemistry A 102, 6601 (1998).

14 A. Ingargiola, R. A. Colyer, D. Kim, F. Panzeri, R. Lin, A. Gulinatti, I. Rech, M. Ghioni, S. Weiss, and X. Michalet, in Proceedings of SPIE, Vol. 8228 (SPIE, San Francisco, CA, USA, 2012) p. 82280B.

15 A. Ingargiola, F. Panzeri, N. Sarkosh, A. Gulinatti, I. Rech, M. Ghioni, S. Weiss, and X. Michalet, in Proceedings of SPIE, Vol. 8590 (San Francisco, CA, USA, 2013) p. 85900E.

16 A. Ingargiola, E. Lerner, S. Chung, F. Panzeri, A. Gulinatti, I. Rech, M. Ghioni, S. Weiss, and X. Michalet, PLOS ONE 12, e0175766 (2017).

17 A. N. Kapanidis, N. K. Lee, T. A. Laurence, S. Doose, E. Margeat, and S. Weiss, Proceedings of the National Academy of Sciences of the United States of America 101, 8936 (2004).

18 A. N. Kapanidis, T. A. Laurence, N. K. Lee, E. Margeat, X. Kong, and S. Weiss, Accounts of Chemical Research 38, 523 (2005).

19 T. A. Laurence, X. Kong, M. Jäger, and S. Weiss, Proceedings of the National Academy of Sciences of the United States of America 102, 17348 (2005).

20 B. K. M. üller, E. Zaychikov, C. Bräuchle, and D. C. Lamb, Bio-physical journal 89, 3508 (2005).

21 N. K. Lee, A. N. Kapanidis, H. R. Koh, Y. Korlann, S. O. Ho, Y. Kim, N. Gassman, S. K. Kim, and S. Weiss, Biophysical Journal 92, 303 (2007).

22 S. W. Yim, T. Kim, T. A. Laurence, S. Partono, D. Kim, Y. Kim, S. Weiss, and A. Reitmair, Clinical Chemistry 58, 707 (2012).

23 S. Doose, M. Heilemann, X. Michalet, S. Weiss, and A. N. Kapanidis, European Biophysics Journal 36, 669 (2007).

24 N. K. Lee, A. N. Kapanidis, Y. Wang, X. Michalet, J. Mukhopadhyay, R. H. Ebright, and S. Weiss, Biophysical Journal 88, 2939 (2005).

25 “Software collection for multispot hardware control,” (), https://github.com/multispot-software.

26 “Software: Repository of data analysis notebooks,” (), https://github.com/tritemio/48-spot-smFRET-PAX-analysis.

27 A. Ingargiola, M. Segal, X. Michalet, and S. Weiss, Figshare (2017), 10.6084/m9.figshare.5146096, DOI: 10.6084/m9.figshare.5146096.

28 A. Ingargiola, T. Laurence, R. Boutelle, S. Weiss, and X. Michalet, Biophysical Journal 110, 26 (2016).

29 A. Gulinatti, I. Rech, P. Maccagnani, and M. Ghioni, in Proceedings of SPIE, Vol. 8631 (SPIE, San Francisco, CA, USA, 2013) p. 86311D.

30 X. Michalet, A. Ingargiola, R. A. Colyer, G. Scalia, S. Weiss, P. Maccagnani, A. Gulinatti, I. Rech, and M. Ghioni, IEEE Journal of Selected Topics in Quantum Electronics 20, 1 (2014).

31 “Notebook: Dcr analysis,” (), http://nbviewer.jupyter.org/github/tritemio/48-spot-smFRET-PAX-analysis/blob/master/DCRplots.ipynb.

32 R. A. Colyer, G. Scalia, I. Rech, A. Gulinatti, M. Ghioni, S. Cova, S. Weiss, and X. Michalet, Biomedical Optics Express 1, 1408 (2010).

33 M. Kloster-Landsberg, G. Herbomel, I. Wang, J. Derouard, C. Vourc’h, Y. Usson, C. Souchier, and A. Delon, Biophysical Journal 103, 1110 (2012).

34 “Notebooks for multispot pattern alignment,” (), https://github.com/tritemio/48-spot-smFRET-PAX-analysis/tree/master/alignment.

35 E. Lerner, S. Chung, B. L. Allen, S. Wang, J. Lee, S. W. Lu, L. W. Grimaud, A. Ingargiola, X. Michalet, Y. Alhadid, S. Borukhov, T. R. Strick, D. J. Taatjes, and S. Weiss, Proceedings of the National Academy of Sciences 113, E6562 (2016).

36 A. Ingargiola, E. Lerner, S. Chung, S. Weiss, and X. Michalet, PLOS ONE 11, e0160716 (2016).

37 “Notebook: multispot 12-bp measurement analysis,” (), http://nbviewer.jupyter.org/github/tritemio/48-spot-smFRET-PAX-analysis/blob/master/smFRET-PAX_single_pop-2017-05-23_08_12d.ipynb.

38 “Notebook: single-spot alex 12 bp analysis,” (), http://nbviewer.jupyter.org/github/tritemio/48-spot-smFRET-PAX-analysis/blob/master/us-ALEX_analysis-2017-06-11_002_12d.ipynb.

39 H. Li, L. Ying, J. J. Green, S. Balasubramanian, and D. Klen-erman, Analytical Chemistry 75, 1664 (2003).

40 “Notebook: multispot 12-22 bp mixture analysis,” (), http://nbviewer.jupyter.org/github/tritemio/48-spot-smFRET-PAX-analysis/blob/master/smFRET-PAX_single_pop-2017-07-11_12d-22d-mixture.ipynb.

41 “Notebook: Multispot vs single spot comparison,” (), http://nbviewer.jupyter.org/github/tritemio/48-spot-smFRET-PAX-analysis/blob/master/smFRET-PAX_ALEX-PAX_comparison_2017-08-03_03_12d.ipynb.

42 R. a. Colyer, G. Scalia, F. a. Villa, F. Guerrieri, S. Tisa, F. Zappa, S. Cova, S. Weiss, and X. Michalet, in Proceedings of SPIE, Vol. 7905 (SPIE, San Francisco, CA, USA, 2011) pp. 790503–790503–8.

43 S. Burri, F. Powolny, C. Bruschini, X. Michalet, F. Regazzoni, and E. Charbon, in Proceedings of SPIE, Vol. 9141 (SPIE, Brussels, Belgium, 2014) p. 914109.

44 J. Buchholz, J. W. Krieger, G. Mocsár, B. Kreith, E. Charbon, G. Vámosi, U. Kebschull, and J. Langowski, Optics Express 20, 17767 (2012).

45 M. Kloster-Landsberg, D. Tyndall, I. Wang, R. Walker, J. Richardson, R. Henderson, and A. Delon, Review of Scientific Instruments 84, 076105 (2013).

46 A. Kufcsák, A. Erdogan, R. Walker, K. Ehrlich, M. Tanner, A. Megia-Fernandez, E. Scholefield, P. Emanuel, K. Dhaliwal, M. Bradley, R. K. Henderson, and N. Krstajić, Optics Express 25, 11103 (2017).

47 C. Bruschini, H. Homulle, and E. Charbon, in Proceedings of SPIE, Vol. 10069 (SPIE, San Francisco, CA, USA, 2017) p. 100691S.

48 A. Ingargiola, P. Peronio, E. Lerner, A. Gulinatti, I. Rech, M. Ghioni, S. Weiss, and X. Michalet, in Proceedings of SPIE, Vol. 10071 (SPIE, San Francisco, CA, USA, 2017) p. 100710Q.

49 A. Ingargiola, T. Laurence, R. Boutelle, S. Weiss, and X. Michalet, in Proceedings of SPIE, Vol. 9714 (SPIE, San Francisco, CA, USA, 2016) p. 971405.

50 D. Tyndall, R. Walker, K. Nguyen, R. Galland, J. Gao, I. Wang, M. Kloster, A. Delon, and R. Henderson, in Proceedings of SPIE, Vol. 8086 (2011) pp. 80860S–80860S–6.

51 A. Ingargiola, bioRxiv, 083287 (2017).

52 “Notebook: multispot 22-bp measurement analysis,” (), http://nbviewer.jupyter.org/github/tritemio/48-spot-smFRET-PAX-analysis/blob/master/smFRET-PAX_single_pop-2017-05-23_04_22d.ipynb.

